# Functional asymmetry and chemical reactivity of CsoR family persulfide sensors

**DOI:** 10.1101/2021.07.25.453692

**Authors:** Joseph N. Fakhoury, Yifan Zhang, Katherine A. Edmonds, Mauro Bringas, Justin L. Luebke, Giovanni Gonzalez-Gutierrez, Daiana A. Capdevila, David P. Giedroc

## Abstract

CstR is a persulfide-sensing member of the functionally diverse copper-sensitive operon repressor (CsoR) superfamily that regulates the bacterial response to hydrogen sulfide (H_2_S) and more oxidized reactive sulfur species (RSS) in Gram-positive pathogens. A cysteine thiol pair on CstR reacts with RSS to form a mixture of interprotomer di-, tri- and tetrasulfide crosslinks, which drives transcriptional derepression of CstR-regulated genes. In some bacteria, notably methicillin-resistant *Staphylococcus aureus* (MRSA), CstR and CsoR, a Cu(I)-sensor, exhibit no regulatory crosstalk in cells, despite maintaining an identical pair of cysteines. We report a sequence similarity network (SSN) analysis of the entire CsoR superfamily, together with the first crystallographic structure of a CstR protein and mass spectrometry-based kinetic profiling experiments to obtain new insights into the molecular basis of RSS specificity in CstRs. The more N-terminal cysteine is the attacking Cys in CstR and is far more nucleophilic than in a CsoR. This cysteine, C30 in *Sp*CstR, is separated from the resolving thiol, C59’, by an Asn55’ wedge. Chemical reactivity experiments reveal a striking asymmetry of reactivity, preserved in all CstRs and with all oxidants tested; however, the distribution of crosslinked products varies markedly among CstRs. Substitution of N55 with Ala in *Sp*CstR significantly impacts the distribution of species, despite adopting the same structure as the parent repressor. We show that CstRs react with hydrogen peroxide, a finding that contrasts sharply with other structurally distinct persulfide sensors from Gram-negative bacteria. This suggests that other factors may enhance the specificity and repressor activity of CstRs in cells.

---

Bacterial pathogens persist in the hostile environment of the infected host as a result of adaptation to host-derived stressors and tolerance to broad-spectrum antibiotics that are used to clear infections. Two broadly recognized anti-bacterial host strategies are the induction of oxidative stress, mediated by myriad highly reactive, inorganic oxygen- and nitrogen species (ROS and RNS), and copper (Cu) toxicity.^1–2^ Bacteria upregulate the biogenesis of hydrogen sulfide (H_2_S) in response to ROS- and antibiotic stress, which enhances persistence in biofilms or in cells^3–4^, while adaptation to Cu poisoning involves cytoplasmic Cu(I) sensing by specialized Cu(I)-sensing metalloregulatory proteins.^5–7^ Cu(I) sensing turns on a transcriptional program to sequester and/or efflux Cu(I) from the cytoplasm of cells. In order for bacteria to deploy H_2_S and H_2_S-derived more oxidized reactive sulfur species (RSS) as infection-relevant antioxidants,^8–9^ cellular H_2_S/RSS levels must be tightly regulated^10^ to avoid H_2_S poisoning and elevated inorganic RSS which can be bactericidal at high levels.^11–12^ RSS-sensing transcriptional regulators have evolved to regulate the expression of RSS biogenesis and clearance enzymes known or projected to reduce cellular loads of H_2_S, RSS and proteome persulfidation, central to the concept of bacterial sulfide/RSS homeostasis.^13–18^

We lack a comprehensive understanding of the chemical specificity and scope of allosteric mechanisms in what we now know are ubiquitous RSS sensors that tightly regulate cellular H_2_S/RSS levels. For example, it is not clear what prevents crosstalk of RSS with metallosensors in cells and the extent to which RSS sensors exhibit specificity over other oxidants, with the possible exception of the RSS-sensing repressor SqrR,^19^ is poorly understood. Detailed investigations of paralogs in the copper-sensitive operon repressor (CsoR) superfamily of bacterial transcriptional repressors^6, 20–21^ present an excellent opportunity to elucidate determinants of biological specificity of closely related proteins.^22–23^ The founding structurally characterized member of the CsoR family is the Cu(I) sensor, copper-sensitive operon repressor (CsoR), initially characterized in *Mycobacterium tuberculosis*.^24^ Since that time, several crystallographic structures have become available for CsoR family repressors,^25^ and each is characterized by a disc-shaped *D*_2_-symmetric or pseudosymmetric tetrameric architecture consisting of ≈90-residue protomers, and exhibit significant pairwise sequence similarities. The CsoR family is known to detect a wide variety of transition metal- and small molecule inducers and comprises other Cu(I) sensors,^26^ including a second mycobacterial Cu(I) sensor, RicR^27^, as well as the Ni(II)/Co(II) sensor RncR from *E. coli*,^28^ the Ni(II) sensor InrS from *Synechocystis*,^29^ the formaldehyde-sensing repressor FrmR from *E. coli*,^22–23^ and the persulfide sensor CstR, now characterized in *Staphylococcus aureus* and *Enterococcus faecalis*.^13–14, 30^ Inducer specificity by CsoR-family repressors was originally described in the context of a simple “W-X-Y-Z” motif using the *Mt*CsoR Cu(I) chelate as a template,^24^ where the first coordination shell ligands around a subunit-bridging, trigonally coordinated Cu(I) define the X (Cys36), Y (His61’) and Z (Cys65’) residue positions.^20^ While insightful, this simple perspective that four residues fully define inducer specificity across a large protein family, engaging in both distinct coordination chemistries and complex cysteine thiol chemistry, has clear limitations.

CsoR and CstR from *Staphylococcus aureus* are CsoR-family paralogs that control distinct physiological processes in the same cytoplasm.^30^ These proteins share 35% sequence identity and 65% similarity, and each contains two conserved cysteine residues in the X and Z positions. *Sa*CsoR recognizes and coordinates Cu(I) with high affinity (*K*_Cu_=10^18.1^ M^-1^) through the two cysteine residues, C41 (C36 in *Mt*CsoR) and C70 (C65 in *Mt*CsoR), and the Nδ1 atom of His66 (H61 in *Mt*CsoR) in a trigonal S_2_N coordination geometry.^24, 30–31^ This results in a decrease in DNA binding affinity, leading to transcriptional derepression of the copper efflux genes *copA*, a P_1B_-type ATPase Cu(I) efflux pump, and *copZ*, a Cu(I) chaperone.^24, 30^ In contrast, *S. aureus* CstR regulates the expression of the *cst* operon that encodes enzymes responsible for mediating resistance to sulfide toxicity and reacts with the sulfane sulfur of both inorganic and organic per- and polysulfides sulfide donors*, e.g.,* glutathione persulfide (GSSH).^13^ Although it is known that CstR, like CsoR, binds Cu(I), Cu(I) binding does not drive dissociation of CstR from *cst* operator DNA *in vitro* or *in vivo*,^13, 30^ as it is missing residues in the second coordination shell (a nearly invariant Tyr35; Glu81 in *Mt*CsoR) previously shown to be energetically linked to Cu(I)-mediated allosteric inhibition of DNA binding.^24, 31^ Moreover, the Cu(I) affinity is such that the intracellular Cu(I) concentration is unlikely to rise to a level required to be bound by CstR, due to intracellular regulation by CsoR^31–32^. This low affinity of CstR for Cu(I), explained by the absence of the Cu(I)-ligating histidine in the Y-position, while helping to explain why CstR is insensitive to Cu(I) in cells, provides no insight into reactant specificity of CstRs or why CsoR fails to sense RSS or other reactive electrophile species in the cell despite of harboring a virtually identical dithiol pair.

In this work, we use a comprehensive sequence similarity network (SSN) analysis of >32,000 sequences to identify three distinct clusters of RSS-sensitive regulators. We validate this analysis by establishing that *S. pneumoniae* D39 SPD_0073 is a *bona fide* CstR and present the first structure of any CstR, which adopts the expected dimer-of-dimers architecture characteristic of the CsoR superfamily.^21, 24^ We find however that while both CsoR^26, 33^ and CstR proteins have long intersubunit S^γ^–S^γ^ distances, a critical determinant in RSS sensing specificity is a far more nucleophilic cysteine in CstRs. Unexpectedly, the two dithiol-based sensing sites on each dimer (C30, C59’) within the tetramer are not equivalent, as evidenced by a significantly faster rate of formation of a singly disulfide-crosslinked dimer vs. the doubly disulfide crosslinked conformer as revealed by comprehensive mass-spectrometry based, kinetically resolved chemical reactivity profiling experiments. This asymmetry characterizes three SSN cluster 10 CstRs, and occurs with both RSS and simple oxidants, but is largely lost in a “wedge” residue substitution mutant (N55A) despite virtually no impact on the structure of the thiol-reduced CstR. We conclude that the rigidity of the dithiol site permits fine-tuning of the reactivity of dithiol pair that may ultimately dictate the sensor specificity for different cellular oxidants and reactive sulfur species in cells.^19^

## Results

### Sequence similarity network (SSN) analysis of CsoR-family repressors delineates distinct functional groups

Early work revealed that founding members of the CsoR family, *M. tuberculosis* copper-sensing repressor (CsoR)^24^ and nickel-sensing repressor *E. coli* RcnR^20, 28^ were representative of a large superfamily from which CstR ultimately emerged, initially characterized in *S. aureus*.^13, 30^ In order to understand the sequence relationships between individual members of this very large superfamily of repressors,^21^ we performed a comprehensive sequence similarity network (SSN) analysis^34^ of 32,476 unique PF02582 entries (11,726 sequences that are <90% identical over 80% of the sequence; Table S1) found in the UniProt database (Fig. 1A) with sequence logo representations of SSN cluster sequence conservation^35^ shown in Fig. 1B. We note that individual SSN clusters represent largely isofunctional groups, as revealed by common sets of regulated genes in the genomic context, as well as by conservation of inducer recognition residues. These functional assignments are further supported by at least one functionally characterized member in each cluster. (Fig. S1). However, several large SSN clusters (6-9; 9.5% of all PF02582 sequences) remain entirely uncharacterized (Fig. S2). Nearly all functionally characterized *bona fide* CsoR-like Cu sensors with the prominent exception of founding member *Mt*CsoR^24^, are grouped in cluster 1 (9450 sequences), which represents the most diverse group, with *Sl*CsoR,^26^ *Mt*RicR,^27^ *Sa*CsoR^30^ and *B*sCsoR^6, 36^ found in distinct subclusters within SSN cluster 1. Despite this diversity, all subclusters conserve the three cognate Cu binding residues that form a C-H-C (Cys-His-Cys) “X-Y-Z” motif and lack an N-terminal conserved metal binding residue “W” (x-C-H-C, Fig. 1). This motif is also present in cluster 23, defined by *Mt*CsoR and found only in actinomycetes, and is characterized by a long, poorly conserved C-terminal tail not found in other CsoRs (Fig. 1B). Another cluster of Cu sensors is represented by *T. thermophilus* CsoR (cluster 17) where one of the Cu(I)- coordinating Cys is replaced by His, defining a x-C-H-H motif (Fig. 1B).^37^ The assignment of SSN cluster 1, 17 and 23 proteins as *bona fide* Cu(I) sensors is further strongly supported by a genomic neighborhood network (GNN) analysis of these sequences, which in the vast majority of cases encodes a CopA efflux pump and a CopZ metallochaperone in the immediate vicinity of the *csoR* gene (Fig. S1).

**Figure 1.**
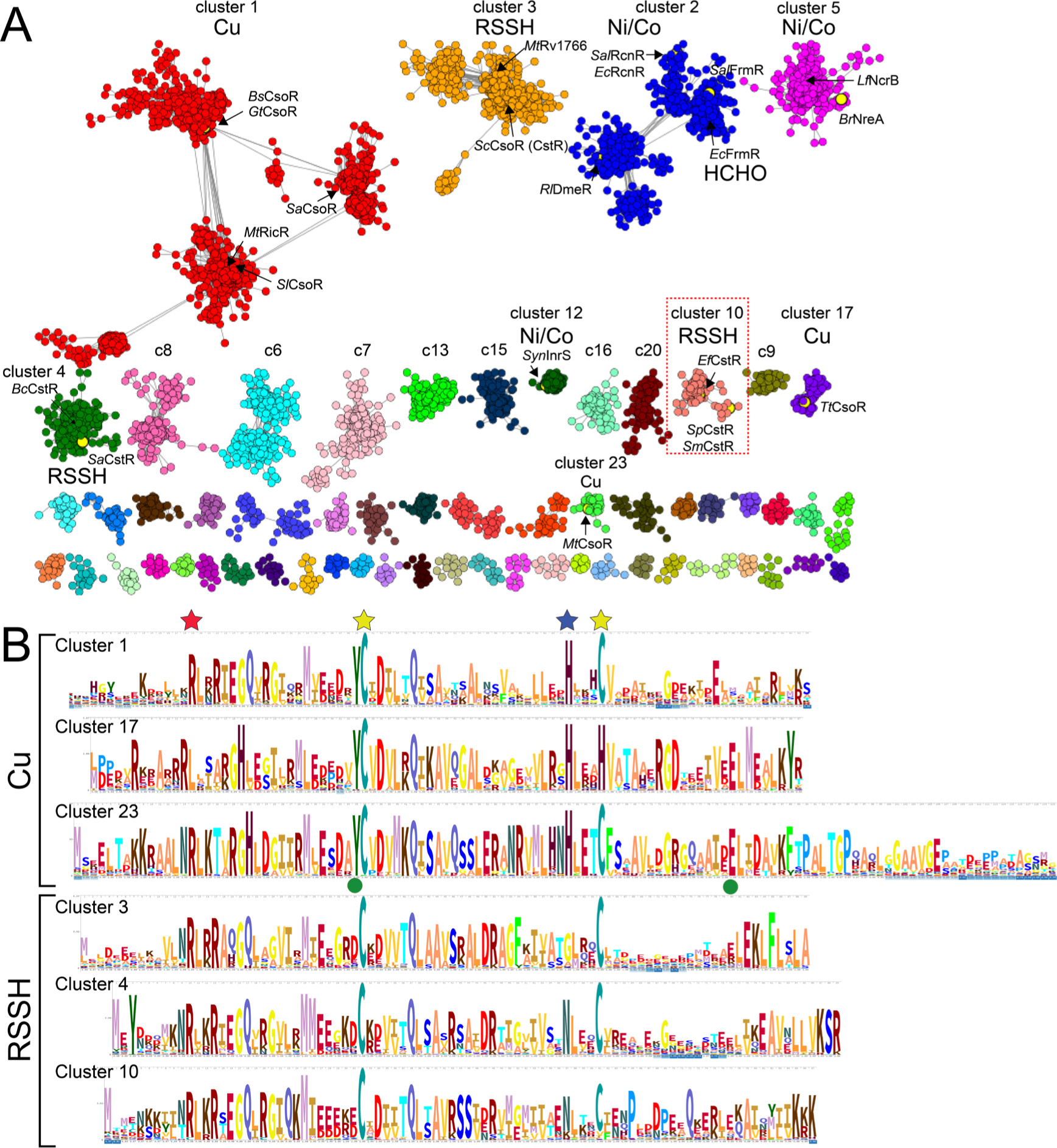
Sequence similarity network (SSN) analysis of the CsoR superfamily of bacterial repressors. (A) Results of an SSN clustering analysis of 32,476 unique sequences belonging to the Pfam PF02582 using genomic enzymology tools and visualized using Cytoscape. Clusters are designated by a number and ranked according to the number of unique sequences (Table S1), color-coded, and functionally annotated as copper (Cu), nickel/cobalt (Ni/Co), persulfide (RSSH) or formaldehyde (HCHO) sensors, if known. Each node corresponds to sequences that are 80% identical over 80% of the sequence, using an alignment score of 28 (see Methods). Functionally characterized members in each of nine clusters (1-5, 10, 12, 17, 23) are indicated with species name and trivial name.^25^ Cluster 10 proteins are the subject of the work presented here. (B) Sequence logo representations of sequence conservation of the Cu(I) and RSSH sensors defined by the indicated cluster of sequences derived from panel A. The residues that coordinate the Cu(I) in CsoRs are marked by the *yellow* and *blue* stars, the residue equivalent to C9 in *Sp*CstR (cluster 10) marked by a *red* star, and the two allosterically important second Cu(I) coordination shell residues in *Mt*CsoR (cluster 23)^87^ marked with a *green* circle. WebLogos for the Ni/Co and other uncharacterized PF02582 sensor clusters are also provided (Fig. S2A).

**Table 1.** Summary of DNA binding parameters for *Sp*CstR and *Sm*CstR.^a^

| Protein | treatment<sup>b</sup> | DNA operator | $K_{\text{tet}}$ (M<sup>-1</sup>) |
| --- | --- | --- | --- |
| C9A *SpCstR* | none | *S. mitis cst* | $2.2 \pm 0.2 \times 10^8$ |
| *SmCstR* | none | *S. mitis cst* | $1.4 \pm 0.2 \times 10^8$ |
| C9A *SpCstR* | none | *Sp adcA* | $4.9 \pm 0.1 \times 10^5$ |
| C9A *SpCstR* | tetrathionate | *S. mitis cst* | $<3.0 \times 10^5$ |
| *SmCstR* | CysSSH | *S. mitis cst* | $<6.3 \times 10^5$ |

<sup>a</sup>Conditions: 10 nM *cst* operator dsDNA in 25 mM Tris-HCl, 200 mM NaCl, 2 mM EDTA, 2 mM TCEP, pH 8.0, 25.0 °C. The DNA binding data were fit to a two-tetramer DNA binding model (see Materials and Methods) and reflect the results of triplicate titrations. <sup>b</sup>none, dithiol-reduced protein.

The second largest cluster of sequences is cluster 2 (6010 unique sequences) and groups the Ni/Co-sensing repressor RcnR, characterized by a H-C-H-H W-X-Y-Z motif, and the formaldehyde (HCHO) sensor, FrmR^23, 38^, characterized by a P-C-H-x W-X-Y-Z motif, on opposite ends of the cluster (Fig. 1 and Fig. S2). The fact that these sequences fail to segregate in our SSN analysis is consistent with a high level of pairwise similarity and the relative ease with which inducer specificity can be relaxed or switched in an FrmR by simply reintroducing the His residue that characterize the RcnR subcluster^23^ (Fig. S2). Here, the GNN analysis can be used to predict the function of individual cluster 2 sequences, as an RcnR or FrmR operon contains either a *rcnA* or an *fmrA* gene, respectively (Fig. S1). Clusters 5 and 12 harbor distinct classes of Ni/Co sensors with a H-C-H-C W-X-Y-Z motif, represented by NreA from *Bradyrhizobium* and *Sphingobium* spp^39^ and NcrB from *Leptospirillum ferriphilum*^40^ (1836 members) and cyanobacterial InrS^29, 41^ (372 members), respectively (Fig. S2). Cluster 3 corresponds to a significant group (3175 members) of largely uncharacterized candidate persulfide (RSSH) sensors from actinomycetes, including *S. coelicolor* CsoR (CstR)^42^ and an uncharacterized mycobacterial protein, *Mt*Rv1766.^24^ Cluster 4 harbors the biochemically characterized RSSH sensor from *S. aureus*,^13^ while cluster 10 contains *E. faecalis* CstR^14^. All the three candidate RSSH sensor clusters (3, 4 and 10) harbor two conserved Cys, forming a x-C-x-C W-X-Y-Z motif and their functional role is confirmed by a genomic neighborhood that encodes for distinct sulfide or sulfite detoxification pathways previously described (Fig. S1)^10, 14^. Overall, our SSN and GNN analyses suggest that the differences in CsoR protein function and inducer recognition can be traced to sequence differences beyond known or projected metal binding residues in the context of the W-X-Y-Z motif model^20–21^, and makes compelling functional predictions for as yet uncharacterized CsoR-family SSN clusters (Fig. S2).

### The nucleophilicity of the N-terminal Cys dictates RSS selectivity over metal ions

Our SSN analysis allows us to define three Cu(I)-sensing clusters and three RSS-sensing clusters of CsoR-family proteins. Clusters 1, 3, 4 and 23 all share two invariant cysteine residues irrespective of their cognate inducers (Cu or RSS) (Fig. S1). We next wished to leverage this analysis to evaluate if the difference in cysteine reactivity (nucleophilicity) could explain the inducer selectivity of the dithiol pair for either Cu(I) ions or RSS. We first performed pulsed alkylation mass spectrometry experiments^43–44^ with the neutral alkylating reagent, *N*-ethylmaleimide (NEM) at pH 7.0 under anaerobic conditions to directly compare the chemical reactivities of the N-terminal Cys in CstR relative to a Cu(I)-sensing CsoR isolated from the same organism, *Staphylococcus aureus* (*Sa*) (clusters 1 vs.4; Fig. 1B). Here, each protein was reacted with a thiol-excess of deuterium-labeled “heavy” NEM (*d*_5_-NEM, pulse) for time, *t*, when an aliquot is removed and added to a denaturing buffer with a vast excess of “light” NEM (H_5_-NEM, chase) to alkylate any unreacted cysteine residues. Proteins were then digested with trypsin and analyzed by MALDI-TOF mass spectrometry (Fig. 2). All observed and expected *d*_5_- and H_5_-NEM derivatized peptide masses are shown in Table S2, with the data modeled by a single or double exponential function (Table S3).

**Figure 2.**
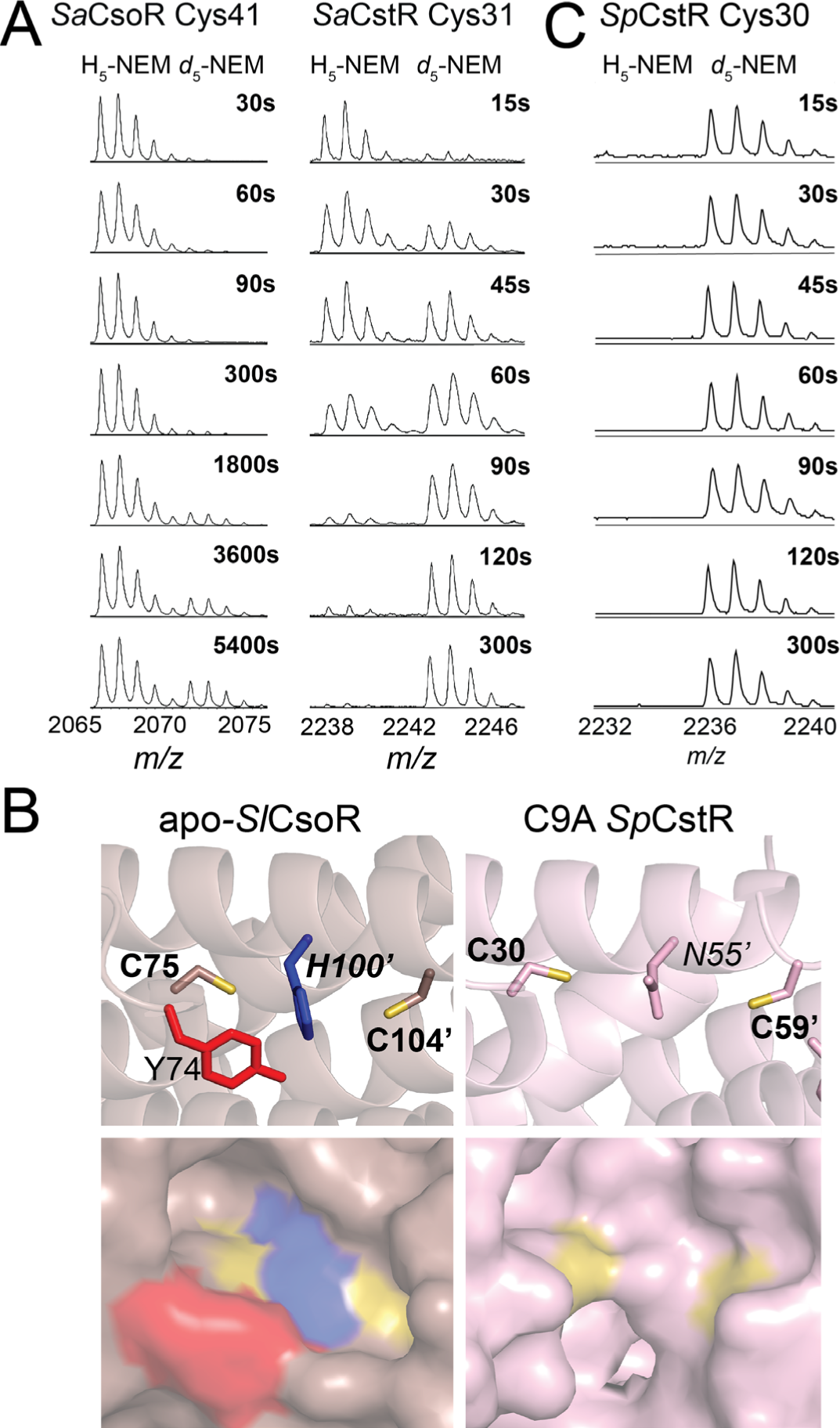
Selected regions of a series of MALDI-TOF mass spectra obtained by rPA-MS of (A) wild-type *Sa*CsoR (*left*) and *Sa*CstR (*right*) showing only the peptide containing the more N-terminal Cys in each case. The *d*_5_-NEM pulse time is indicated on each panel, with the H_5_-NEM and *d*_5_-NEM isotopic mass distributions shown, with expected and measured monoisotopic masses^67^ shown for each tryptic peptide shown compiled in Table S2. Representative MALDI-TOF spectral regions for Cys9, Cy30 and Cys59 of *Sp*CstR are provided (Fig. S3). The data in panel were analyzed using single or double exponential model (see Fig. S3) with the kinetic parameters compiled in Table S3. (B) Structures of the dithiol regulatory sites shown for the Cu(I) sensor *Streptomyces lividans* (*Sl*) CsoR in the apo-state^33^ (*left*) and the persulfide C9A SpCstR (*right*) (this work). Residues of interest are shown. (C) Selected region of a series of MALDI-TOF mass spectra obtained by rPA-MS of wild-type *Sp*CstR.

**Table 2.**
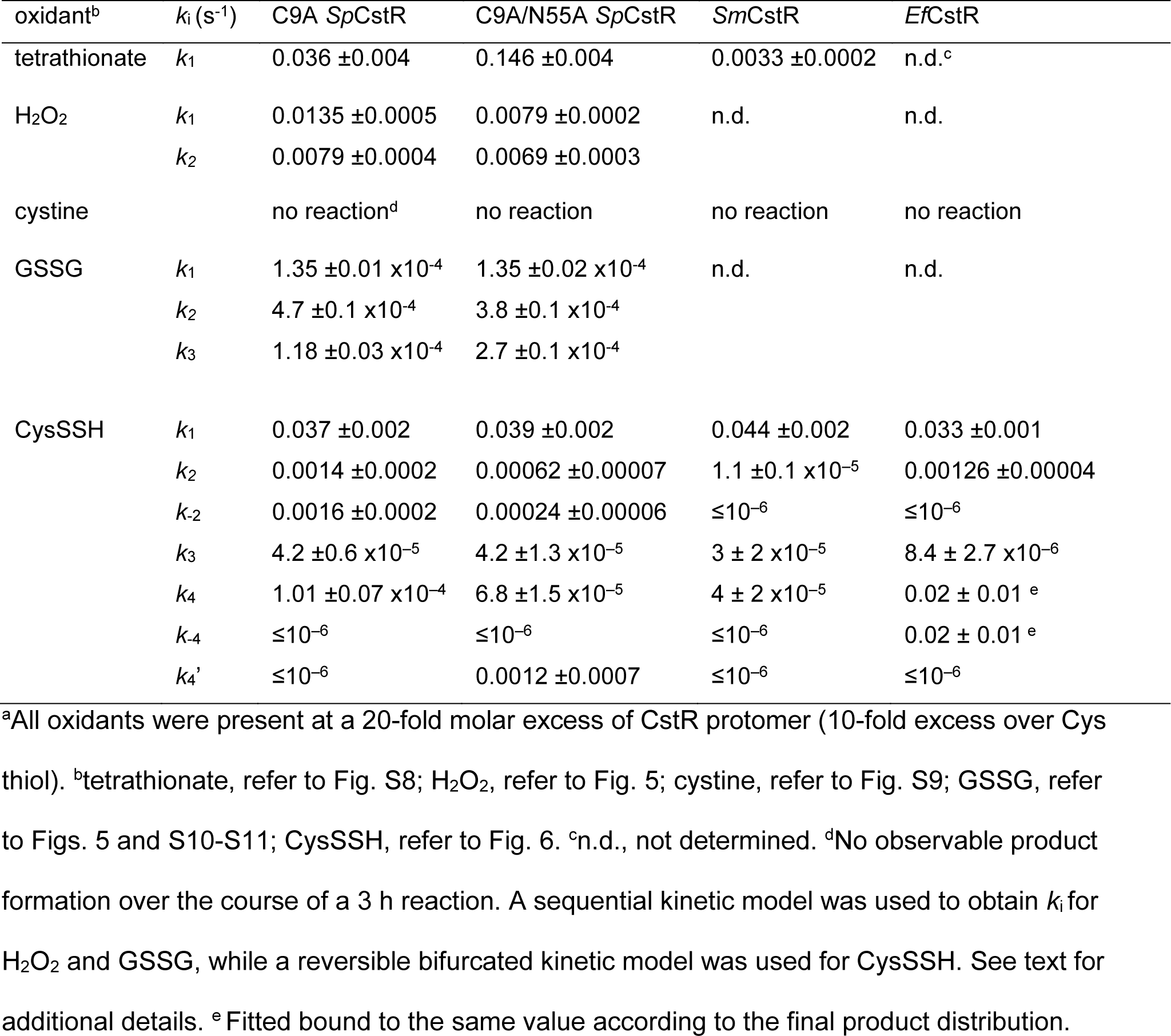
Compilation of all rate constants observed in the thiol kinetic profiling experiments with the indicated oxidant^a^

We find that the reactivity of the more N-terminal Cys in *Sa*CsoR (C41) is very slow (Fig. 2A) when compared to the more C-terminal Cys (C70) (Fig. S3), which is characterized by a nucleophilicity largely as expected for a solvent-exposed thiolate. Indeed, inspection of a structure of apo-CsoR from *S. lividans* reveals that while C104 (C70 in *Sa*CsoR) is on the edge of a tetramer in a solvent exposed helix, C75 (C41 in *Sa*CsoR) is boxed into a cage-like structure formed by the Cu-ligating His100’ (H66’ in *Sa*CsoR; analogous to N55 in *Sp*CstR) and the key allosteric residue Y74 (Y40 in *Sa*CsoR), that are conserved among all Cu(I) sensing clusters (Fig. 1 and Fig. 2B, *left*).^31^ This is also the case for *Tt*CsoR in the apo-state, where C41 is found in the bottom of a cavity, while C66 is located on a solvent-exposed loop that is not resolved in this structure. These observations suggest that a restriction of solvent accessibility by the N-terminal cysteine attenuates its reactivity and thus enforces Cu(I) specificity.

In striking contrast, in *Sa*CstR the N-terminal Cys (C31) is *more* reactive than the C-terminal Cys (C60) (Fig. 2A; Table S3) and a comparison of reactivities of the N-terminal Cys in *Sa*CstR vs. *Sa*CsoR reveals a 50-fold difference in the second order rate constant, a trend that persists over a wide range of pH (6.0-9.5) (Fig. S3). Indeed, these data suggest that Cys31 in *Sa*CstR has a p*K*_a_ of ≈8.6, or similar to that of a free thiol, 8.5-9.0,^45–46^ while Cys41 in *Sa*CsoR is elevated to ≈9.3 or higher, consistent with its decreased reactivity over the entire pH range (Fig. S3). Furthermore, this relative trend of cysteine nucleophilicity is common among other RSS-sensors found in cluster 10, including the previously characterized *Ef*CstR^14^ and two candidate CstRs from Streptococcal *spp* (*vide infra*). Cluster 10 CstRs (Table S3) exhibit an even higher the reactivity of the N-terminal Cys relative to *Sa*CstR (Fig. 2C), with *Sm*CstR nearly too fast to measure. These findings reveal that the microenvironment near C41 in CsoR makes deprotonation less favorable than in a CstR, thus providing a mechanistic rationale for a lack of transcriptional regulatory crosstalk between CstR and CsoR sensors in the *S. aureus* cytoplasm.^30^ Moreover, these findings make the prediction that the N-terminal cysteine of a CstR is solvent-accessible relative to the same residue in a Cu(I)-sensing protein.

### Streptococcus pneumoniae D39 SPD_0073 is a CstR

In order to identify new CstR proteins that would serve as structural targets to test the predictions from our pulsed alkylation experiments, we took advantage of our SSN analysis to make functional predictions for individual protein nodes contained within a single SSN cluster.^47^ We turned our attention to a single cluster 10 node, encoded by a gene designated locus tag SPD_0073 in the serotype 2 pneumococcal strain D39.^48^ SPD_0073 possesses significant sequence similarity to the previously described CstR from *E. faecalis* (cluster 10; Fig. 1A) but is more distantly related to *S. aureus* CstR, and not nearby other genes that suggest a role in RSS sensing and metabolism (Fig. 3A). This prompted a much broader search for other cluster 10 candidate CstRs which are closely related to SPD_0073. We identified a candidate CstR encoded in mitis group oral streptococci closely related to *S. pneumoniae*^49^ that is known to encounter fluctuations in H_2_S in periodontal biofilms.^50^ The operon bears strong resemblance to the sulfide- and RSS-inducible *cst*-like operon that we characterized in *E. faecalis*, encoding two single-domain sulfurtransferases or rhodaneses, RhdA and RhdB, and a coenzyme A persulfide reductase (Fig. 3A; Fig. S1).^14^ Those operons are found in most strains of *S. constellatus*, *S. anginosus*, *S. cristatus*, *S. gallolyticus* and *S. mitis*, and also harbor a consensus CstR binding sequence in the operator-promoter region upstream of the *rhdB* gene.^30^ These features make the prediction that these operons would be inducible by Na_2_S or RSS.^14^

**Figure 3.**
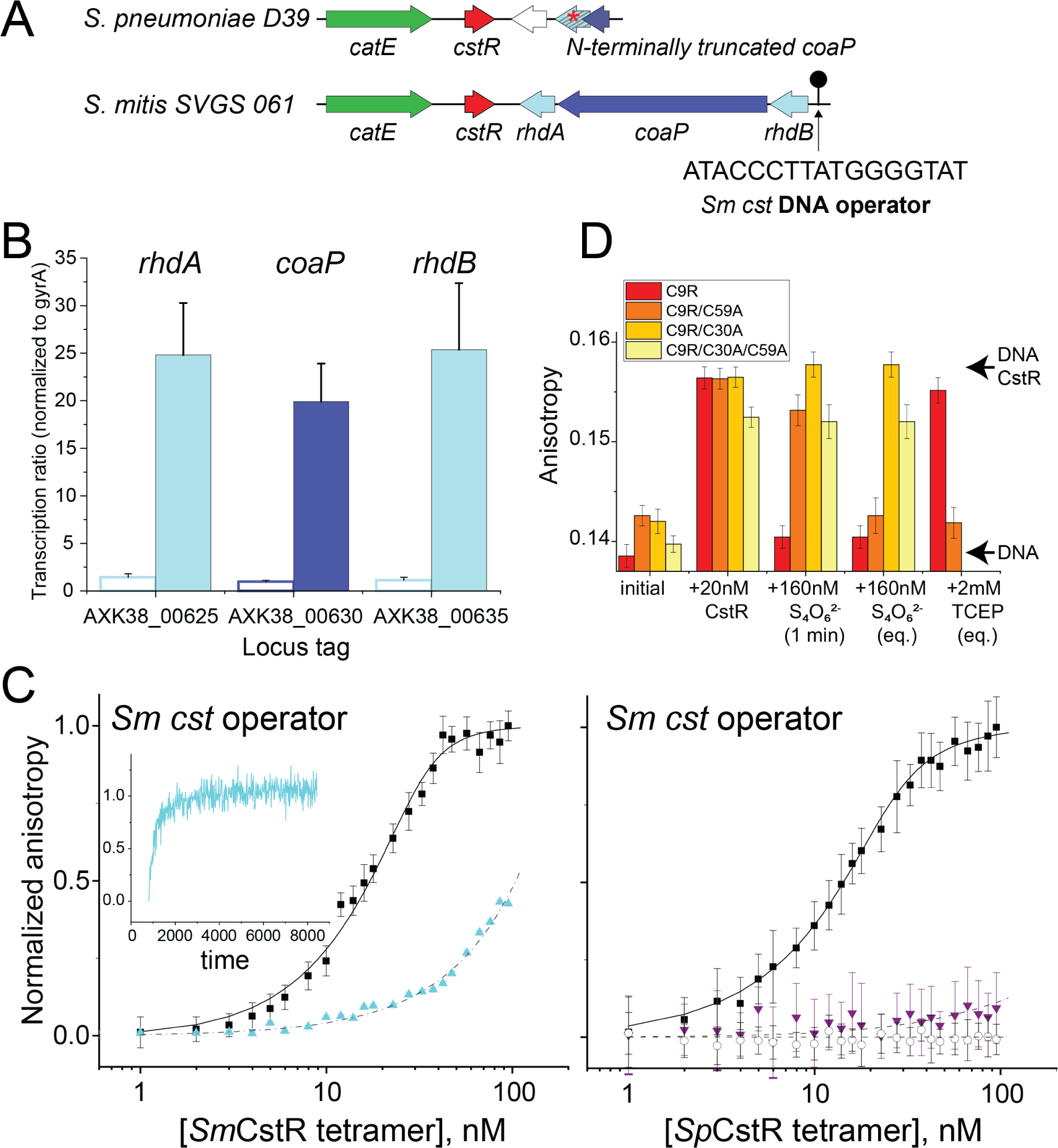
*S. pneumoniae* (*Sp*) D39 SPD_0073 encodes a *bona fide* persulfide-sensing CstR. (A) Genomic neighborhood and location of a candidate CstR-regulated operator-promoter site around selected streptococcal *cstR* genes (cluster 10; Fig. 1A) investigated here. The DNA sequence of the core *Sm cst* operator is shown.^30^ (B) Induction of the expression of genes in *S. mitis* SVGS_061 by exogenous sulfide (*shaded* bars; *white* bars, no treatment) as measured by qRT-PCR. AXK38_00625 (*rhdA*; see panel A); AXK38_00630 (*coaP)*; AXK38_00635 (*rhdB*). (C) *Left*, *Sm*CstR (locus tag AXK38_00620) binds to the *S. mitis* SVGS061 *cst* operator-containing DNA (*black* circles) and is inhibited by pre-treatment with CysSSH (*cyan* triangles) which is reversible upon addition of TCEP (*cyan* time course, *inset*). *Right*, representative DNA binding curves obtained for reduced (*black* circles) or tetrathionate-treated (*purple* triangles) C9A *Sp*CstR on the *S. mitis* SVGS061 *cst* operator-containing DNA. *Black* open circles, C9A *Sp*CstR binding to an AdcR operator,^88^ representative of non-specific binding. The continuous lines drawn through each data set represent the results of a fit to a two-tetramer binding model (Fig. S3)^30^, with parameters complied in Table 1. (D) Bar chart representation of the kinetics of *Sm cst* operator binding, dissociation by tetrathionate and subsequent reduction by TCEP by various *Sp*CstR dithiol site mutants in a C9R *Sp*CstR parent background, from *left* to *right*. The arrows labeled DNA CstR and DNA are the anisotropies associated with the CstR-bound and free DNAs. 1 min, anisotropy 1 min after addition of tetrathionate; eq, equilibrium anisotropy value.

To test this prediction, we focused on *S. mitis* SVGS_061, isolated from the bloodstream of an infected cancer patient.^51^ The *S. mitis* CstR (AXK38_00620) is 95% identical to SPD_0073 (Fig. S4) and a comparison of the genomic neighborhoods around putative *cstR*-encoded genes suggests a precise genomic deletion in D39, relative to the *S. mitis* strain (Fig. 3A).^48^ We recovered this strain and assessed induction of the candidate CstR-regulated genes (AXK38_00625, *rhdA*; AXK38_00630; *coaP*; AXK38_00635, *rhdB*) by growing cultures on a rich growth medium (brain-heart infusion, BHI) with early mid-log cells treated with 0.2 mM Na_2_S, prior to collecting RNA 30 min post-induction (Fig. 3B). We observe robust sulfide-inducible expression of candidate CstR-regulated genes, establishing *S. mitis* CstR as a *bona fide* persulfide sensor. We confirmed the sulfide-induced regulation by studying DNA binding affinities of purified wild-type *S. mitis* CstR and the C9A *S. pneumoniae* SPD_0073 (this mutation has no impact on DNA binding and was introduced to avoid undesirable oxidative chemistry at this non-conserved Cys; Fig. S4A). We carried out quantitative fluorescence anisotropy-based DNA binding experiments with a duplex harboring the *S. mitis cst* operator and analyzed with a two-tetramer binding model (Fig. 3A; Fig. S4B).^36, 52^ These experiments show that the average tetramer binding affinity (*K*_tet_) is virtually identical for both proteins (Fig. 3C; Table 1) and is significantly weakened in a *reversible* fashion upon incubation with RSS or a reactive disulfide-inducing electrophile, tetrathionate (Fig. 3C-D), provided that C30, and to lesser degree C59, is present. A van’t Hoff analysis of the DNA-binding thermodynamics reveals that assembly of two CstR tetramers on the DNA operator is largely driven by a favorable enthalpy change, consistent with previous studies of a *Streptomyces* ssp. CsoR^52^ and a modest electrostatic component (Fig. S4C-D). These studies establish that *S. pneumoniae* SPD_0073 exhibits DNA binding properties and regulation consistent with that of a CstR, which we denote *Sp*CstR.

### The crystallographic structure of C9A *Sp*CstR reveals a more accessible N-terminal Cys and functional asymmetry that is confirmed by cysteine reactivity experiments

Having validated the SSN-predicted functions of *Sp*CstR and *Sm*CstR, we turned to crystallography to further understand the structural basis of RSS-sensing specificity. We solved the crystallographic structure of reduced C9A *Sp*CstR at pH 7.5 from 34% PEG200 to 2.3 Å resolution. This first crystal structure of a CstR shows the anticipated dimer-of-dimers pseudo-tetrameric architecture (Fig. 4A; Table S4 for structure statistics), which each protomer harboring three helices (α1-α3). This is compatible with our small angle x-ray scattering (SAXS) and NMR spectroscopy analysis that indicates that *Sp*CstR adopts a homogeneous tetrameric assembly state at pH 7.5 (Fig. S5 & S6). Despite the relatively low resolution, the structure reveals a microenvironment around the N-terminal Cys that is far less sterically hindered when compared to structures of apo-CsoR (Fig. 2B). These structural data are fully compatible with the chemical reactivity data above (Fig. 2C).

**Figure 4.**
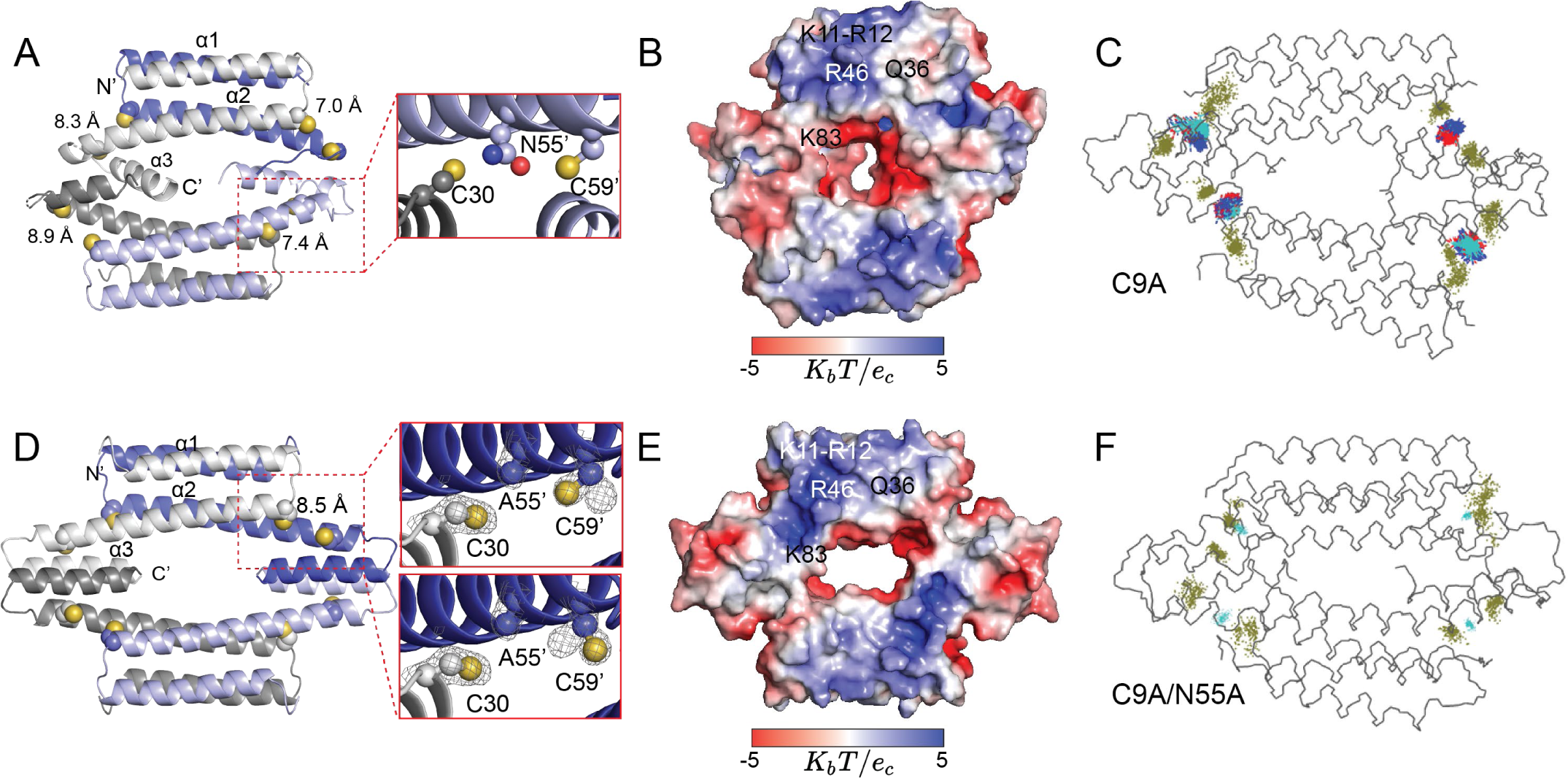
Ribbon and electrostatic surface representations of the crystallographic structures of homotetrameric C9A *Sp*CstR (A, B, respectively) at pH 7.5 and C9A/N55A *Sp*CstR (D, E), at pH 6.0. See Table S2 for structure statistics. In panels A and D, each of the protomers are shaded *white*, *blue*, *grey* and *light blue*. The side chains of C30, N55 (A55) and C59 are presented in sphere representation, with CPK coloring and S^γ^ atoms *yellow*. Interprotomer C30 S^γ^-C59’ S^γ^ distances are indicated. In C9A/N55A *Sp*CstR, the S^γ^ atom of C59’ occupies two rotamer positions. (C) and (F), results of molecular dynamics (MD) simulations obtained for C9A *Sp*CstR derived from *in silico* mutation of C9A/N55A *Sp*CstR to C9A *Sp*CstR as the starting structure (C) or authentic C9A/N55A *Sp*CstR as starting structure (F). The temporal displacements of the Cys side chains (*yellow*) and A55 (C) or N55 (F) over the course of these simulations are shown.

The *Sp*CstR tetramer is highly asymmetric, with an asymmetric unit of 1 protomer, with one dimer laterally offset relative to the other, and a wide range of C30 S^γ^-C59’ S^γ^ distances, but with the side chain of N55 generally “wedged” between the two Cys side chains, maintaining long S^γ^–S^γ^ interprotomer distances (7.0 to 8.4 Å) (Fig. 4A). This observation suggests that long S^γ^–S^γ^ interprotomer distances are a common feature of dithiol CsoR family proteins irrespective of their cognate inducer (Fig. 2C). Surprisingly, the orientations of the C-terminal α3 helices in each protomer at the tetramer interface are distinct in this model (Fig. S5A). Small angle x-ray scattering (SAXS) analysis reveals that *Sp*CstR adopts a homogeneous tetrameric assembly state at pH 7.5 (Fig. S5B), while NMR spectroscopy shows that any deviations from a *D*_2_-symmetric tetrameric structure at this pH must occur on a fast timescale, since only one set of peaks is observed in the ^1^H-^15^N TROSY spectra (Fig. S6, right panel). Moreover, an additional structure solved at pH 5.5 to a resolution of 2.0 Å (Table S4) adopts a distinct assembly state in the crystal lattice, accommodated by significant displacements of the α3 helix (Fig. S5C, *top*). The difficulties obtaining a high-resolution structure of the tetrameric assembly are also reflected in our molecular dynamics simulations starting with the C9A *Sp*CstR at pH 7.5, where the interface between protomers has significant mobility and does not necessarily reach an equilibrium state (Fig. S8A). These data suggest that packing of the dimer-dimer interface is somewhat plastic in C9A *Sp*CstR.

In order to explore the functional impact of the structural asymmetry on chemical reactivity, we implemented a mass spectrometry-based kinetic profiling approach. Pseudo-first order anaerobic reactions were carried out in at least triplicate with 20-fold molar excess oxidant over protomer, quenched with excess iodoacetamide (IAM, +57 amu) at time *t*, and proteins were subjected to ESI-MS. While tetrathionate reacts quantitatively within the first ≈1 min with no evidence of intermediates (Fig. S9), hydrogen peroxide (H_2_O_2_) and glutathione disulfide (GSSG) show three and four major species, respectively (Fig. 5A; Table S5). In addition to the two end-states, fully reduced/capped monomer (*black*) and doubly disulfide-crosslinked dimer (denoted *di/di*; *blue*), we observe a transiently populated, on-pathway intermediate dimer species in which one side of the dimer is closed with a disulfide, with the other side remaining reduced (or sulfenylated^53^), and subject to capping by IAM, or in the case of GSSG, singly- or doubly-glutathionylated (Fig. 5B). The differences in the rate of formation of the first disulphide bond in the dimer unit relative to the formation of the second at the other side of the dimer unit constitutes direct evidence for asymmetry of reactivity in the *Sp*CstR dimer-of-dimers architecture. This observation is consistent with our crystal structure that appears to have trapped an asymmetric state that is otherwise interconverting rapidly in solution by NMR spectroscopy (Fig. S7). Furthermore, given the long distance between the two thiols in the crystal, disulfide closing of one dithiol site may introduce strain on the opposite site within the dimer that inhibits the subsequent closure as a result of this asymmetry.

**Figure 5.**
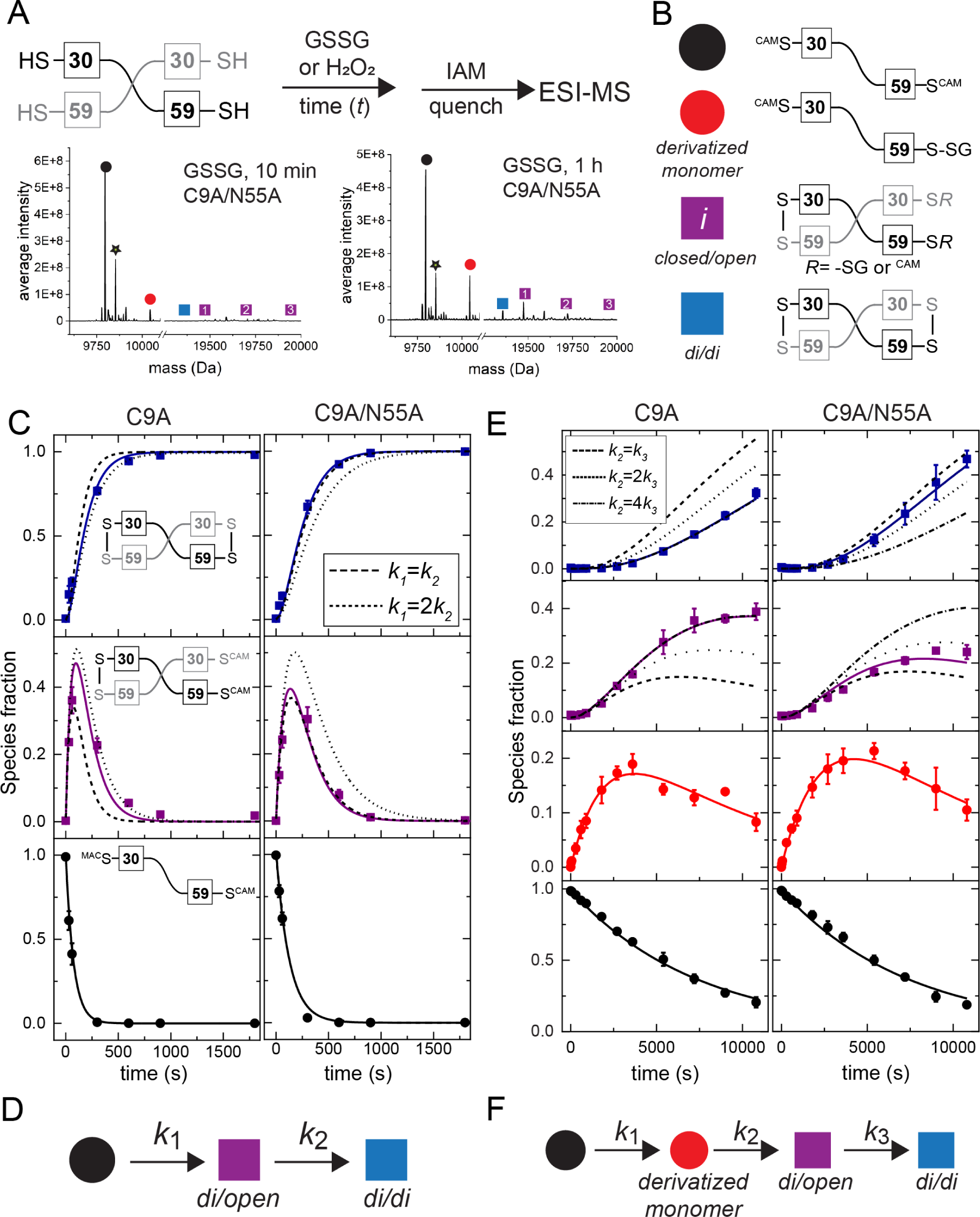
Thiol reactivity kinetic profiling experiments with non-sulfur-based oxidants, hydrogen peroxide (H_2_O_2_) and glutathione disulfide. (A) (*top*) Overall workflow with a schematic non-covalent “dimer” representation of *Sp*CstR shown with the Cys residues indicated. (*bottom*) Representative regions of the ESI-MS data obtained for a 10 min (*left*) or 60 min (*right*) reaction of GSSG with C9A/N55A *Sp*CstR. The symbols correspond to the state harboring distinct modifications on one or both thiols as shown in panel (B). Tables S5-S6 lists expected and observed masses for each of these species. (C) Representative kinetic profiling experiments obtained for C9A *Sp*CstR (*left*) and C9A/N55A *Sp*CstR (*right*) and H_2_O_2_. (D) Kinetic scheme used to analyze these data with *k*_i_ compiled in Table 2, and defined by the continuous lines (*dashed* line, symmetric limit, *k*_1_= *k*_2_, and *dotted* line, two-fold asymmetry, *k*_1_=2*k*_2_, in panel (C). (E) Representative kinetic profiling experiments obtained for C9A *Sp*CstR (*left*) and C9A/N55A *Sp*CstR (*right*) and GSSG. (F) Kinetic scheme used to analyze these data with *k*_i_ are compiled in Table 2, and defined by the continuous lines (*dashed* line, symmetric limit, *k*_1_= *k*_2_; *dotted* line, two-fold asymmetry, *k*_1_=2*k*_2_, and *dashed-dotted* line four-fold asymmetry, *k*_1_=4*k*_2_, in panel E.

### An N55A substitution diminishes functional asymmetry in *Sp*CstRs

Our results thus far suggest that *Sp*CstR harnesses a highly reactive N-terminal attacking Cys capable of forming a disulfide with a resolving C-terminal Cys in which the S^γ^ atoms are separated by 7.0 to 8.4 Å; formation of that disulfide, in turn, further enhances the asymmetry observed in the *Sp*CstR crystal structure that inhibits, to some extent, the formation of a symmetrically disulfide-crosslinked (*di/di*) dimer. We reasoned that N55, which replaces the Cu(I)-coordinating His in Cu(I)-sensing CsoRs (Fig. 1B), might enforce this asymmetry of reactivity, given its positioning as a “wedge” residue between the cysteines. To test this, we solved the crystallographic structure of C9A/N55A *Sp*CstR to 1.4 Å at pH 6.0 (Table S4) and investigated its kinetics of reactivity toward various oxidants. The structure reveals a highly symmetric tetrameric architecture (Fig. 4D), in striking contrast to the parent C9A *Sp*CstR structure, confirmed as a tetrameric assembly state by SAXS analysis (Fig. S5D). Interestingly all the other structural features remain unaltered relative to C9A *Sp*CstR. Long S^γ^–S^γ^ interprotomer distances (8.6 Å) in two rotamer populations of C59 are observed, as is an unbroken α2 helix, despite the loss of the N55 side chain. Furthermore, sequential backbone (^1^H_N_, ^13^Cα, ^13^C’ and ^15^N_N_) NMR resonance assignments for both C9A and C9A/N55A *Sp*CstRs at pH 5.5 reveal that the N55A substitution gives rise to only the anticipated local perturbation in the α2 helix, which propagates to the structurally adjacent region of the C-terminal α3 helix (Fig. S7A). The TALOS-derived secondary structures of the two variants are identical and fully consistent with C9A/N55A *Sp*CstR structure (Fig. S7B).

We next turned to our mass spectrometry-based kinetic profiling to interrogate the effect of this N55A cavity mutation on chemical reactivity and functional asymmetry. We find that the N55A substitution reduces the effect of asymmetry in chemical reactivity with both GSSG and H_2_O_2_ (Fig. 5, Table 2). Consistently, molecular dynamics simulations of C9A/N55A *Sp*CstR reveal a highly symmetric conformational ensemble (Fig. 4F, Fig. S8B). Moreover, an *in silico* A55-to-N55 substitution reintroduces asymmetry at the level of the two dithiol sites on opposite ends of the tetramer within in dimer, as reflected by the low spread of visited coordinates of the S^γ^ atoms in half of the sites (Fig. 4C, F). These C9A *Sp*CstR simulations are remarkably reminiscent of the structure of the formaldehyde sensor FrmR with formaldehyde adducts at half of the dithiol sites (Fig. 4C, F).^22^ Furthermore, these MD simulations are consistent with our NMR dynamics that show lower motional disorder in C9A *Sp*CstR compared to the C9A/N55A mutant in the linker that connects the α2 and α3 helices although the α1-α2 linker changes less (Fig. S7). We propose that even if the asymmetry of C9A *Sp*CstR is of greater amplitude at longer timescales, the mechanical strain imposed by the presence of N55 defines two sets of thiol conformations interconverting on a timescale compatible with the temporal resolution of the NMR and SAXS experiments. Overall, these MD simulations as well as the NMR and SAXS data from C9A and C9A/N55A *Sp*CstRs support the idea that functional asymmetry arises from fast internal dynamics in the loops that may impact packing in the dithiol sensing sites on opposite sides of the *Sp*CstR tetramer, an extreme example of which is simply trapped in our highly asymmetric structure of C9A *Sp*CstR (Fig. 4A).

### Relative stability of crosslinked products upon reaction with thiol persulfides varies among cluster 10 CstRs

We next wished to determine if these specific features of structural and functional asymmetry observed with *Sp*CstR can be extended to other SSN cluster 10 RSS-sensing repressors and to other biologically relevant oxidants, *e.g*., cysteine persulfide found in cells (CysSSH) (Fig. 6).^8, 14, 18, 54^ CysSSH is generated *in situ* by anaerobic incubation of cystine with an excess Na_2_S, forming equimolar CysSH and CysSSH, with remaining HS^−^ and cystine, the latter of which does not react with CstR thiols (Fig. S9) and is used immediately. Previous evidence suggests that thiol persulfides react rapidly with CstRs, but are known to form a mixture of di-, tri- and tetrasulfide linkages, at least when incubated for long periods of time.^13^ Further, in at least one case, the major oxidative product in a CstR from the actinomycete *Streptomyces coelicolor* (cluster 3, Fig. 1A) is reported to be a per- or polysulfide on the N-terminal Cys.^42^ We therefore implemented a mass spectrometry-based kinetic profiling approach (Fig. 6A) to elucidate the mechanism and rates of product formation for three cluster 10 CstRs (*Sp*CstRs, *Sm*CstR and *E. faecalis* CstR) toward CysSSH, in an effort to understand if the asymmetry observed in *Sp*CstR plays a role in fine-tunning thiol reactivity towards the cognate inducer.^19^

**Figure 6.**
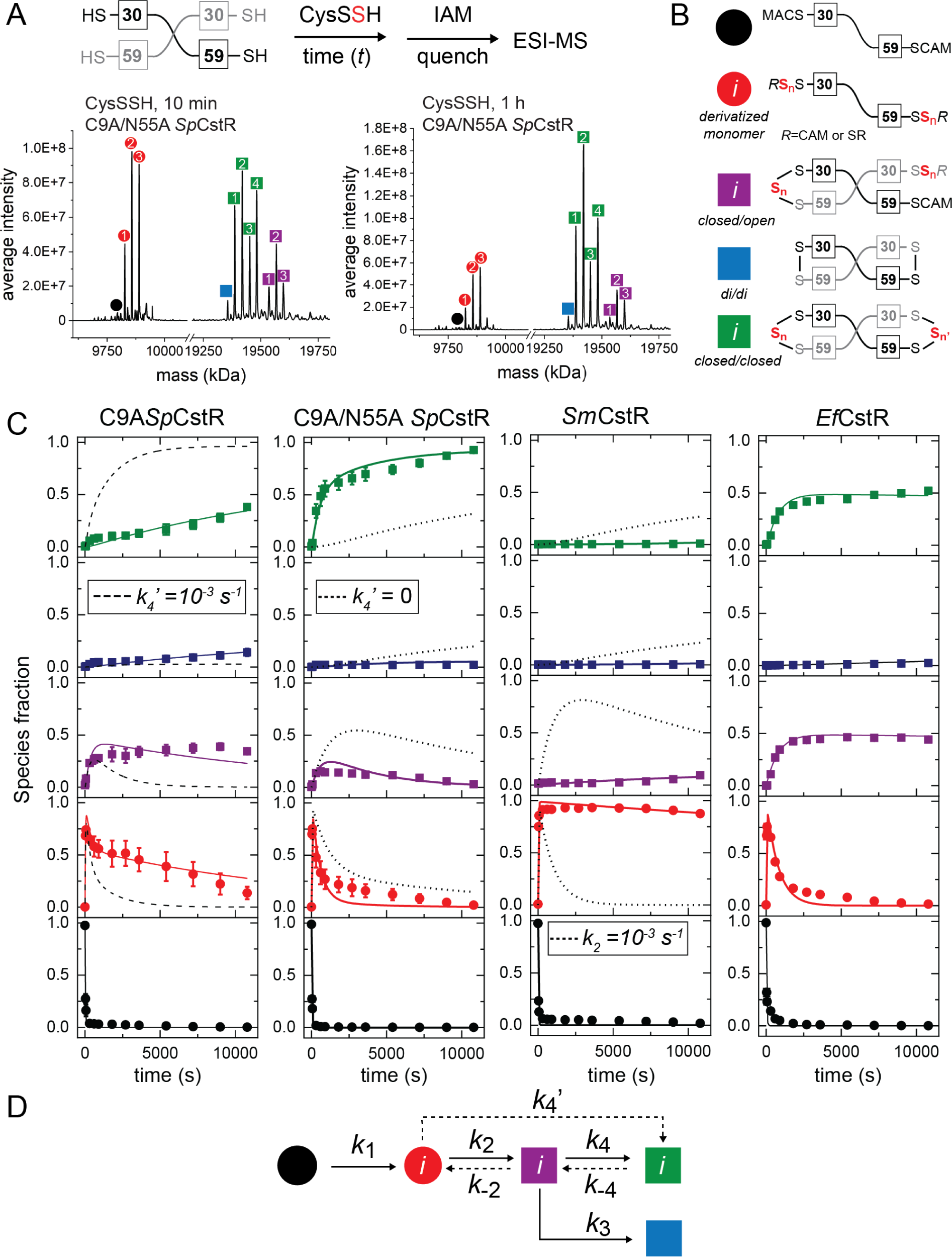
Cys persulfide (CysSSH) reactivity kinetic profiling experiments with cluster 10 CstRs. (A) Overall workflow. A schematic noncovalent “dimer” representation of *Sp*CstR or *Sm*CstR (one-half of the tetramer) is shown, with the Cys residues indicated. For *Ef*CstR, Cys residues corresponding to C30 and C59 in *Sp*CstR are C32 and C61, respectively. Representative regions of the ESI-MS data obtained for a 10 min (B) or 60 min (C) reaction of CysSSH with C9A/N55A *Sp*CstR. In each case, the monomer region is shown at *left*, and the crosslinked dimer region is shown at *right*. The symbols correspond to the generalized structures shown in panel D, with numbers within each symbol corresponding to the *i*th state harboring *n* distinct +32 species distributed among one or both thiols (isomers cannot be resolved using this method): *i*=1, *n*=1; *i*=2, *n*=2; *i*=3, *n*=3; *i*=4, *n*=4. Table S7 lists expected and observed masses for each of these species. (C) Global analysis of these kinetic profiles obtained for C9A *Sp*CstR, C9A/N55A *Sp*CstR, *Sm*CstR and *Ef*CstR (*left* to *right*) with CysSSH, using the generalized kinetic scheme in panel D. The species are colored according to the key shown in panel B, with the sum of all *i*th species grouped into each data point. The continuous lines drawn through each dataset reflect the results of global fitting to the minimal bifurcated model shown in panel D, with individual *k*_i_ compiled in Table 2. (D) Generalized kinetic scheme used to analyze the reaction profiles obtained with persulfide donors. Continuous lines represent the processes that were determined in all profiles, whereas dashed lines represent the processes that could only be determined for some of the proteins. Rate constants are compiled in Table 2.

As expected from previous work, the dithiol persulfide-sensing site in CstRs is capable of forming a variety of protomer crosslinked oxidized products upon persulfide treatment in a way that depends on the incubation time (Fig. 6A).^13^ Five groups of reaction products are needed to fully describe the kinetic data (Fig. 6B, Table S7) and were fit to a kinetic scheme in which a thiol-derivatized per- and polysulfide protomeric (*red*) intermediate containing up to three sulfur additions capped by IAM and comprising more than 70% of species at early time points, is irreversibly formed from the reduced and capped monomer (*black*), defined by *k*_1_ (Fig. 6D). This gives rise to three groups of covalently crosslinked species, *closed*/open (*purple*), a di/di (*blue*) species introduced above, and a new *closed/closed* (*green*) dimer (see Fig. 6B), which harbor C30-C59’ tri- or tetrasulfide crosslinks (Fig. 6C, *green*).^13^ The kinetic data for C9A *Sp*CstR was globally fitted using a linear model that incorporates the tri- or tetrasulfide crosslinks as another *closed*/*closed* (*green*) product from the *closed*/*open* intermediate (*purple*, Fig. 6C, *left*). Also, since the derivatized monomer is not completely consumed, a rate constant for its formation from the *closed*/*open* intermediate was introduced (*k_-2_*, Table 2). Although the formation of *closed/closed* products with tri- and tetrasulfide linkages are favored in C9A *Sp*CstR with respect to di/di species (*k_4_ > k_3_*), the formation of these species is slow. This is not unexpected given the kinetic barrier to formation of the symmetric di/di product with other oxidants (Fig. 5). In contrast, treatment of C9A/N55A *Sp*CstR with CysSSH yields nearly quantitative formation of *closed*/*closed* product (*green*) in a time-dependence that is best modelled with the formation of these species from the polysulfidated monomer (*red*) with no accumulation of *closed*/*open* intermediates (*purple*). Notably, numerical simulations with different rate constants of formation of *closed*/*closed* product from the polysulfidated monomer (*k_4_’*, Table 2, Fig. 6C, *dashed* and *dotted* lines) can explain the differences in the kinetic profiles between C9A and C9A/N55A *Sp*CstRs nearly entirely. This observation suggests that the loss of the bulky N55 side chain removes a kinetic block for the formation of higher order tri- and tetrasulfide crosslinked dimers within the CstR tetramer, supporting the idea that asymmetry determines not only the kinetic profile but also the final product distribution.

Remarkably, the reactivity profile observed for *Sm*CstR collapses to one highly populated (≈90%) species, the polysulfidated monomer (*red*), with successive sulfur additions (up to three) presumably added to C30 (Fig. 6C, Table S7). A very small amount of asymmetric di-open species is formed at long time points, with the “open” site again harboring mixed disulfide linkages to cysteine. We see no evidence for the formation of di/di, di/tri and other *closed*/*closed* species over a 3 h reaction. Analysis using the same kinetic scheme (Fig. 6D) reveals a significant decrease in the rate constant for the formation of linkages between the C30 and C59 (*k_2_*, Table 2, Fig. 6C, *dotted* lines). These kinetic profiling data with *Sm*CstR appear to represent an extreme case of the reactivity profile of C9A *Sp*CstR (Fig. 6C) except that no *closed/closed* species are formed. It is interesting to note that this product distribution resembles that obtained previously with a cluster 3 CstR.^42^ This kinetic profiling data is compatible with a more significant kinetic block for the formation of higher order tri- and tetrasulfide crosslinked dimers within the CstR tetramer for *Sm*CstR. Indeed, the *Sm*CstR tetrathionate reactivity profile confirms that this protein can access a *closed*/*closed* conformation but does so significantly more slowly relative to *Sp*CstRs (Fig. S10; Table 2). It is interesting to note that these differences in product distributions originate from just four amino acid substitutions, one of which is the non-conserved C9 in *Sp*CstR (Fig. S4). Of the remaining three amino acid differences between *Sm*CstR and *Sp*CstR, G26, derived from a non-conservative E26G substitution, is most conspicuous (Fig. S3A). G26 terminates the α1 helix in *Sp*CstR and defines a point of increased backbone motional disorder on the sub-ns timescale by NMR (Fig. S7). This Gly is positioned above the attacking Cys, C30, and thus may markedly impact its range of chemical reactivity in comparing these two closely related CstRs (Fig. 4A). Indeed, MD simulations of an *in silico* G26E substitution mutant reveal that E26 engages in several salt bridges with neighboring residues that may inhibit formation of di- and polysulfide crosslinks in *Sm*CstR (Fig. S8C).

The reactivity profile of *Ef*CstR, on the other hand, bears close resemblance to that of C9A/N55A *Sp*CstR except that the relative concentrations of the *closed*/*open* species is higher, accumulating to ≈40% of all species (*purple*, Fig. 6C), with correspondingly less *closed*/*closed* species formed (*green*, Fig. 6C). This kinetic profile can be modelled considering a fast equilibrium between *closed*/*open* and *closed/closed* species (defined by *k_4_* and *k_-4_*) (*red*). This fast interconversion and the relatively high kinetic constants suggest that *Ef*CstR is even more flexible and less kinetically trapped than C9A/N55A *Sp*CstR. This observation also suggests that the E26 present in both *Sm*CstR (kinetically trapped) and *Ef*CstR (fast interconversion from open to closed) is not necessarily sufficient to create a kinetic block, and other residues differences between *Sm*CstR and *Ef*CstR might be responsible for the dramatic change in reactivity (Fig. S4). Altogether our kinetic profiling experiments with CysSSH suggest that functional asymmetry and kinetic barriers for the formation of linkages can be tuned with minimal sequence alterations that significantly impact the product distribution.

## Discussion

In this work, we present new insights into the structure and chemical reactivity of an organic persulfide sensor derived from the CsoR superfamily of bacterial repressors.^10, 13^ We have defined the structure and sequence features that distinguish a Cu(I)-sensing CsoR from a persulfide-sensing CstR, focusing on characterized proteins from *S. aureus*, a human pathogen known to encode both of these dithiol CsoRs with no evidence of crosstalk in cells.^30^ Our SSN and GNN analyses allow us to identify three clusters of putative persulfide sensors, from which we were able to functionally and structurally characterize for the first time a CstR, representative of closely related persulfide sensors from the lactic acid bacteria.^14^ These structures and comprehensive chemical reactivity data suggest the presence of high kinetic barriers toward the formation of symmetrically crosslinked states. Further, this asymmetry can be traced to internal dynamics that can be fine-tuned by minimal sequence perturbations in what is rather limited sequence diversity within a single cluster of protein sequences.

Our structures of *Sp*CstR in the dithiol-reduced and DNA-binding competent state, when placed in the context of conserved sequence features of the cluster 10 family repressors, reveal features that are both common to, and distinct from, Cu(I)-sensing CsoRs of known structure. *Sp*CstR is characterized by a pair of symmetry-related ion pairs across the α1-α1’ dimer interface, E14-R18’, and an additional ion pair in the α1-α2 loop, R28-D32, which surrounds C30, and thus may help orient the attacking thiol in CstR; both features are widespread in the CsoR superfamily.^24^ A prominent stripe of positive electrostatic potential crosses one face of the tetramer, over the positively charged “hole” of the tetramer (R84 and K85 are highly dynamic and not visible in this structure) and includes superfamily-invariant residues, K11 and R12 of the RXXK/R sequence (X, any amino acid) in α1 helix (C9-X-K11-R12 in *Sp*CstR),^24, 52, 55^ Q36^52^ and R46 in the α2 helix and the C-terminal K83-R84-K85 motif,^55–56^ all known to play energetically important roles in DNA binding (Fig. 4). A major point of departure between CstR and CsoR is the intrinsic reactivity of the N-terminal Cys which is far higher in a CstR and is readily explained by the differences in microenvironment in apo-*Sl*CsoR^33^ vs. reduced *Sp*CstR. Indeed, close approach of Cu(I) to the Cu(I) sensing site in CsoR would result in the concerted formation of the two strong Cu(I) coordination bonds to the Cys and His, which upon recruitment of the more distal C-terminal Cys into the first coordination shell, breaks the helical sense of the α2 helix, enforcing an allosteric switch that ultimately drives CsoR off the DNA.^6, 33, 57^

In addition to understanding what distinguishes a Cu(I) sensor from an RSS sensor, the extent to which an RSS sensor is specific for one oxidant (RSS) over another remains the subject of debate.^58–59^ It has recently been shown that *R. capsulatus* SqrR and BigR from an animal pathogen, which adopt entirely distinct structures from that of CstR beyond a common dithiol persulfide sensing site^15–16, 19^ can distinguish RSS from ROS by means of structural “frustration” of the disulfide form.^19^ In striking contrast, CstRs react rapidly with H_2_O_2_, but at a ≈3-5-fold slower rate than observed with CysSSH. This contrasts with what has been observed in *S. aureus*, where it was shown that H_2_O_2_ added to the outside of cells does not induce the *cst* operon.^13^ On the other hand, more recent work reveals that a range of potent oxidants, including polysulfane compounds, are capable of inducing transcription of CstR-regulated genes.^12, 60^ Thus, the persulfide specificity of a CstR may well be lower than for SqrR/BigR, but the primary physiological (vs. pathophysiological) inducer is likely reactive sulfur species.

In this context, it is interesting to speculate on the extent to which a relaxed degree of oxidant specificity may impact bacterial physiology. Cluster 10 proteins are derived nearly exclusively from the order Lactobacillales or lactic acid bacteria that are generally non-respiring and catalase-negative, acid-tolerant, low GC-content Gram-positive bacteria that meet their energy needs via carbohydrate fermentation with the production of lactic acid as a waste product. These organisms lack a TCA cycle and an electron transport chain that can generate significant endogenous ROS. Some *Streptococcus* spp., *e.g., Streptococcus mutans*, are important components of dental plaque, a biofilm that forms on the surface of teeth, and NADH oxidase is capable of reducing any O_2_ to water, regenerating NAD^+^, thus limiting exposure to other reactive oxygen species.^61^ As a result, the apparent inability of CstR to readily distinguish H_2_O_2_ from RSS in these species is unlikely to be physiologically relevant, even in microaerobic conditions. However, this likely is not the case for *Streptococcus pneumoniae*, a facultative anaerobe that is uniquely capable of respiring on oxygen by employing pyruvate oxidase (*spxB*), and to a lesser extent, lactate oxidase (*lctO*). These enzymes collectively convert lactic acid to acetyl-phosphate via pyruvate, to generate copious intracellular H_2_O_2_ that is used to limit microbial competitors during the colonization of the respiratory tract^62–63^ and induce host cell damage in the presence of host-derived nitric oxide.^64^ It is interesting to note that *S. pneumoniae* is unique among cluster 10 organisms in that it has clearly lost the RSS sensing and detoxification response (see Fig. S2) found other lactic acid bacteria, such as *S. mitis* and *E. faecalis*. This is consistent with the idea that RSS would not be expected to accumulate in *S. pneumoniae*, due to high levels of endogenous H_2_O_2_.^8^

A striking finding is that the kinetic barrier to the formation of symmetric di-, tri- or tetrasulfide crosslinks dictates the product distribution of CstRs upon treatment with a biologically relevant persulfide donor, cysteine persulfide. The relative impact of this kinetic barrier appears to be determined by the number of accessible states of the S^γ^ atoms in the dithiol sensing site, which is tunable by minimal sequence changes. The flexibility of this site determines the probability of resolving a polysulfide built on the attacking cysteine and, thus, the relative abundance of species harboring di-, tri- or tetrasulfide linkages. An extreme case is *Sm*CstR, where even long incubations with CysSSH give rise to only small amounts of the di- open species, below 5% (Fig. 6). This finding is consistent with a strong structural asymmetry in a “half-the-sites” reacted dimer. Interestingly, the crystal structure of the CsoR-family formaldehyde-sensing repressor FrmR is characterized by the same tetrameric asymmetry, with one site “closed” by a methylene bridge initiated by Michael addition of formaldehyde to the sole attacking Cys (equivalent to C30 in *Sp*CstR), and the other sensing site in the same dimer unreacted.^22^ No reaction kinetics or product distributions were presented in that work, but this finding is consistent with the kinetic profiling we present here for CstRs and with the asymmetric tetramer structure of C9A CstR at pH 7.5, in which the two dimers are strongly offset relative to one another, in striking contrast to the C9A/N55A *Sp*CstR structure (Fig. 4). Based on these results and those others, we suggest that the microenvironment around each cysteine thiol and the dynamic flexibility of those sites gives rise to a spectrum of derivatized and crosslinked species that is far more complex than anticipated on the basis of other sulfur-nitrogen and sulfur-sulfur bridges found in other allosteric redox switches in regulatory proteins and enzymes.^19, 22, 65–66^ Ongoing studies are directed toward elucidating the persulfide switching mechanism in CstRs, will begin to elucidate to what extent this diversity in thiol reactivity impacts bacterial physiology.

## Materials and Methods

### Sequence Similarity Network (SSN) and Genomic Neighborhood Network (GNN)

The EFI-EST (http://efi.igb.illinois.edu/efi-est/) and EFI-GNT (http://efi.igb.illinois.edu/efi-gnt/) web tools were used to generate SSNs and GNNs, respectively. SSNs were generated using the Option B (Families) option of EFI-EST using the Pfam of *Sp*CstR (PF02582; trans_repr_metal). An alignment score of 28 was used to generate the SSN with no minimum or maximum number of residues, and the final network generated was 80% representative (repnode 80), collapsing sequences of 80% identity over 90% of the sequence into a representative node and was visualized using Cytoscape (http://www.cytoscape.org/). The composite FASTA file containing all unique sequences associated with each cluster of interest was used to generate a multiple sequence alignment (MSA) through Jalview, using ClustalO with default parameters with sequences containing long N-terminal and C-terminal extensions on either side of a core region manually removed to facilitate comparison of sequences. MSAs were then uploaded to Skylign (https://skylign.org/) to generate logo plots associated with each SSN cluster.^35^ The list of sequences used to generate the logo plot are provided in Table S1 arranged in individual clusters with Uniprot IDs associated with biochemically characterized CsoR-family proteins highlighted in each cluster.

### Protein Purification

*Staphylococcus aureus* CsoR and CstR and *Enterococcus faecalis* CstR were purified from His-tagged constructs to homogeneity in a fully reduced form as previously described.^13–14, 30^ *E. coli* expression plasmids encoding 6x-His-tagged wild-type, C9A and C9A/N55A *S. pneumoniae* CstR (locus tag SPD_0073) and S. mitis SVGS_061 CstR (locus tag AXK38_00635) were transformed into BL21(DE3) competent cells and allowed to grow overnight at 37 °C. This preculture was used to inoculate LB containing 100 μg/mL ampicillin and allowed to grow at 37 °C until an OD of 0.6-0.8 was reached. 0.5 mM IPTG was then added, and the growth allowed to continue overnight at 16 °C. Cells were harvested by centrifugation for 20 min, 4 °C (3000 xg), resuspended with 20 – 30 mL/g wet cells of degassed low imidazole buffer (25 mM Tris, 500 mM NaCl, 20 mM imidazole, 10% glycerol, pH 8.0) and sonicated for a total of 5 min with a 2 s on/8 s off pulse sequence. The lysate was clarified by centrifugation (10 min, 17000 xg). Polyethyleneimine (PEI) was added to the supernatant at a final concentration of 0.2% v/v and allowed to stir for 40 min at 4 °C. The resulting fraction was centrifuged (10 min, 17000 xg) and the soluble fraction subjected to ammonium sulfate precipitation (70% w/v) at 4 °C (50 min) and the remaining solution centrifuged (10 min, 17000 xg) and the pellet used for Ni-NTA affinity chromatography. Low imidazole buffer and high imidazole buffer (25 mM Tris, 500mM NaCl, 500 mM imidazole, 10% glycerol, pH 8.0) were prepared and degassed and TCEP was added to 2.0 mM. An imidazole gradient was used to elute CstRs, peak fractions collected, subjected to TEV cleavage and dialyzed, centrifuged (10 min, 13,000 rpm) or filtered through 0.45 μm filter and the flow-through fractions from the Ni-NTA pooled and subjected to size exclusion chromatography (Superdex 75; GE, Boston, MA) in degassed G75 buffer (25 mM Tris, 200 mM NaCl, 2 mM EDTA, pH 8.0) and peak fractions were collected and stored at –80 °C until use. A typical yield was 25 and 30 mg/L culture for *S. pneumoniae* and *S. mitis* CstRs, respectively, and all contained the correct number of reduced cysteines by Ellman’s reagent. Protein concentrations were determined by UV absorption using the following ε_280_ values: 2980 M^-1^ cm^-1^, for *S. pneumoniae*, *S. mitis* and *E. faecalis* CstRs.^14^

### Quantitative RT-PCR analysis sulfide-induced genes in *Streptococcus mitis* SVGS_061

S. mitis SVGS_061 strain was inoculated from glycerol stocks and grown in 5 mL BHI medium overnight. The overnight culture was pelleted by centrifugation, resuspended in equal volume of BHI and diluted into 30 mL BHI medium with starting OD_600_ of 0.01. Triplicate cultures were grown to an OD_600_ of 0.2 aerobically at 37 °C with shaking (200 rpm) in loosely capped 50 mL Falcon tubes, followed by the addition of freshly prepared 0.2 mM Na_2_S. 5 mL aliquots were withdrawn from the cultures prior to addition of Na_2_S (*t*=0), and at 30 min post addition, and centrifuged for 10 min. Cell pellets were washed with ice-cold PBS, re-centrifuged for 5 min, decanted and stored at –80 °C until analysis by qRT-PCR using the primers specific for the *rhdA* (locus tag AXK38_00625), *coaP* (locus tag AXK38_00630) and *rhdB* (locus tag AXK38_00635) genes, essentially as previously described.^13, 18^

### Ratiometric Pulsed-Alkylation Mass Spectrometry (rPA-MS)

Sample preparation for pulsed-alkylation mass spectrometry was adapted from the original report of the technique^43^ and optimized for CstR and CsoR. All experiments were carried out anaerobically in a glovebox in a buffer containing 10 mM HEPES and 200 mM NaCl at pH 7.0. CstR, CstR mutants, and CsoR proteins were reacted with a 3-fold molar thiol excess of *d*_5_-*N*-ethylmaleimide (NEM, pulse, Isotech). At discrete time points, 50 μL aliquots were removed and quenched with an equal volume of a solution containing a 900-fold thiol excess of H_5_-NEM (chase) with 100 mM Tris pH 8.0, and 8 M urea. After a 1 h chase, quenched reactions were removed from the glovebox and precipitated on ice with a final concentration of 12.5% trichloroacetic acid (TCA) for 1.5 h. Precipitated protein was pelleted by centrifugation at 4° C. The supernatant was removed, and the pellet washed twice with ice-cold acetone. The washed pellet was vacuum centrifuged to dryness at 45° C and re-suspended in 10 μL digest buffer (20 mM ammonium bicarbonate, 10% acetonitrile, 50:1 protein:trypsin ratio, pH 8.2). CstRs were digested for 1 h and CsoR for 0.5 h at 37° C. Tryptic digests were quenched with a final concentration of 1% trifluoroacetic acid (TFA) and spotted on a MALDI plate with α-cyano-4-hydroxycinnamic acid (CCA) matrix with a 5:1 matrix:sample ratio for analysis. For examination of the pH-dependence of the reaction, the same experiment was carried out in the following buffers; Mes for pH 6.0-6.5, HEPES for pH 7.0-7.5, and Tris for pH 8.0-9.0.

MALDI-TOF mass spectra were collected and analyzed in triplicate reactions using a Bruker Autoflex III MALDI-TOF mass spectrometer with 200 Hz frequency-tripled Nd:YAG laser (355 nm) and Flex Analysis software (Bruker Daltonics, Billerica, MA). Cysteine-containing peaks were identified by monoisotopic mass (Table S2) and resolved as alkylated with d_5_-NEM (+130.0791 Da) or H_5_-NEM (+125.0477 Da) with little to no detectable unmodified peptide (data not shown). The theoretical distribution and peak areas were determined using the averagine algorithm^67^ and quantified by summing the total peak areas of the full isotopic distribution. Relative peak areas were used to determine the mol fraction of H_5_-NEM labeled peptide, Θ(H_5_), as defined by Equation 1. A(H_5_) and A(d_5_) correspond to the area (A) of the isotopic distribution of H_5_-NEM or d_5_-NEM alkylated peptide, respectively. To obtain the pseudo-first order rate constant of alkylation, *k*, Θ(H_5_) was plotted as a function of pulse time, *t*, and fit to Equation 2. In some instances, a fit to a sum of two exponentials was used, Equation 3. The second-order rate constant was obtained by dividing *k* by the concentration of *d*_5_-NEM in the pulse.

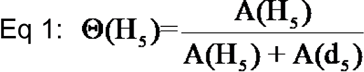

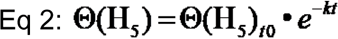

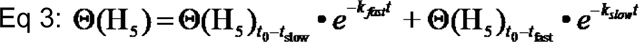

### Fluorescence Anisotropy Titrations

Double-stranded 5’-fluorescein-labeled DNA constructs were annealed as component single-strands and titrated as previously described using an ISS PC1 Spectrofluorometer (Champaign, IL) equipped with an automatic titrator unit.^44^ The *S. mitis cst* operator, 5’-5FluoroTACCTCCAAAT**<u>ATA</u>CCCTTAT-GGGG<u>TAT</u>**ATTAAA and the complementary unlabeled strand, was used for all experiments (core *cst* operator sequence is in bold).^30^ 10 nM *cst* operator dsDNA in 25 mM Tris-HCl, 200 mM NaCl, 2 mM EDTA, 2 mM TCEP, pH 8.0 was titrated with either WT or the indicated CstR mutant (≈12 µM stock). Fluorescein was excited at 495 nm and polarization monitored with a 515 nm cut-off filter in the L-format. Each data point collected was the average and standard deviation of five measurements. Normalized *r* values for the fractional saturation of *cst* OP1 was calculated as (*r*_obs_ – *r*_0_)/(*r*_complex_ – r_0_) from 0 to 1 where *r*_complex_ represents the maximum anisotropy obtained and *r*_0_ represents free *cst* OP1 DNA. For titrations not reaching saturation, *r*_complex_ was calculated from the addition of the anisotropy change of reduced CstR to *r*_0_ of the corresponding non-saturating CstR. Collected data were fit to a sequential non-dissociable tetramer (CstR_4_) binding model^13–14, 30^ using DynaFit^68^ assuming a linear relationship between *r*_obs_ and *v*_i_ and the binding density at the *i*th addition of titrant.^30^ The macroscopic binding constant, *K*_tet_, is reported and was determined as *K*_tet_ = (*K*_1_•*K*_2_) due to high uncertainty in extracting unique *K*_i_ values, *K*_1_ and *K*_2_, due to a strong inverse correlation and little sigmoidal behavior in the binding isotherms as previously described.^30^ Reported affinities (*K*_tet_) are the average of three independent experiments.

### Anaerobic mass spectrometry-based kinetic profiling of CstRs with various oxidants

Protein stocks for these experiments transferred into the argon-filled glove box and buffer exchanged into either 25 mM Tris, 200 mM NaCl, pH 8.0 (for tetrathionate, cystine, H_2_O_2_, glutathione disulfide reactions) or 150 mM sodium phosphate pH 7.4, and 2 mM EDTA (for the *S*-nitrosoacetyl penicillamine (SNAP) donor^69^ and cysteine persulfide) using PD MiniTrap G-25 (GE, Boston, Massachusetts) column, followed by repeated centrifugation (10 min, 13,000 rpm, 6 times). Reaction mixtures contained 60 µM protomer (15 µM tetramer) and a 20-fold molar excess over Cys residues (10-fold over CstR protomer) of each oxidant (1.2 mM) in a total volume of 500 µL Triplicated reactions were performed by withdrawing 50 µL at each time, *t* (ranging from 30 s to 3 h) and mixed with 50 μL of a 900-fold excess of iodoacetamide (IAM). Prior to initiating the reaction with oxidant, 50 μL was withdrawn in the same way, as *t*=0 sample. IAM quenching the reaction and caps (amidomethylates) thiols and per- and polysulfides that form on Cys residues. Quenched reactions were then removed from the glovebox and washed with buffer three times in a 10 kDa mini-concentrator (Millipore-Sigma, Burlington Massachusetts) and 5 uL of each sample was aliquoted into a screwcap vial (Waters, Milford, Massachusetts) with 45 uL of buffer and analyzed by LC-ESI-MS, essentially as previously described.^19^ using a Waters/Micromass LCT Classic time-of-flight (TOF, Milford, MA) mass spectrometer with a CapLC inlet following chromatography on a 50 µm Agilent BioBasic C4 reverse-phase column (5 µm particle size, 300 Å pore size). A 20 min linear gradient from 10% Solvent A (5% acetonitrile, 95% water, 0.1% formic acid) to 90% Solvent B (95% acetonitrile, 5% water, 0.1% formic acid) was used with the elution monitored at 215 nm. Data were collected and analyzed using MassLynx Software (Waters, Milford, MA).

For data analysis, species with peak intensities greater than 20% that of the most intense feature at every timepoint were picked and the intensities for each peak from triplicate runs were averaged and normalized to determine relative abundance. For dimeric species, a factor of 4 was used to account for the decreased ionization efficiency of the dimer relative to monomeric CstR species, obtained by comparing peak intensities of a CstR in the reduced vs. tetrathionate-oxidized *di/di* form prepared under identical conditions. There was no significant impact of the nature of the CstR on this intensity ratio (data not shown). The normalized intensities of each of five species that share defined chemical features and kinetic profiles, shown in Tables S5-S7 (*black* circles, *red* circles, *blue* squares, *green* squares and *purple* squares), were then binned, and a standard calculated, and the resulting progress curves globally analyzed (no weighting function was used) by applying the simplest possible kinetic model, which was a linear model, unless otherwise stated, using Dynafit.^68^ Schematic representations of these fitting models are shown in each case. Data were fitted using a pseudo-first order reaction, with the concentration of oxidant at a 20-fold excess, and kinetic rate constants compiled in Table 2.

### Small Angle X-ray Scattering (SAXS)

Reduced C9A and C9A/N55A CstR samples were prepared to a final concentration of 1 mg/mL, 3 mg/mL, and 5 mg/mL and data acquired at Argonne National Laboratory (Argonne, Chicago, IL) and analyzed by the Small-Angle X-ray Scattering Core Facility, Frederick National Laboratory for Cancer Research, Frederick, MD, according to published procedures.^6^ Briefly, the buffer scattering was subtracted from the sample scattering, with modified sample scattering curves merged to generate the scattering curve for further analysis. The scattering data at low *q* values are used to estimate the radius of gyration (*R*_g_) by the Guinier approximation with the range of *qR*_g_<1.3. Pair distance distribution function (PDDF) plots of each protein were obtained using the distance distribution analysis tool on SAS Data Analysis (Primus). *D*_max_ values were calibrated in order to make the PDDF curve drop smoothly to zero.

### X-ray Crystallography

C9A *Sp*CstR, selenomethionine (SeMet) C9A *Sp*CstR, and C9A/N55A *Sp*CstR (6-8 mg/mL) crystals were grown at 20°C using the hanging-drop vapor-diffusion method. Crystals formed under the following conditions: C9A *Sp*CstR (PDB code 7MQ1): 0.1 M MES, pH 5.5, 0.25 M (NH_4_)_2_SO_4_, 15-18% PEG 4000; C9A SeMet *Sp*CstR (PDB code 7MQ2): 0.1 M HEPES, pH 7.5, 34-40% PEG 200; C9A/N55A *Sp*CstR (PDB code 7MQ3): 0.1 M MES, pH 6.0, 50 mM CaCl_2_, 40-50% PEG 200. Crystals were harvested, cryo-protected in a reservoir solution supplemented with 25% glycerol, except for C9A/N55A *Sp*CstR, and were flash-frozen in liquid nitrogen. Diffraction data were collected at 100 K at the Beamline station 4.2.2 at the Advanced Light Source (Berkeley National Laboratory, CA) and were indexed, integrated, and scaled using XDS. A complete dataset of the native C9A *Sp*CstR in space group C2 at ≈2 Å and a redundant dataset of the same variant at λ=1.77118 nm were collected. Despite the excellent anomalous signal for sulfur in this dataset, phase estimation using Single Anomalous Diffraction (SAD) was unsuccessful, as were several heavy metal soaks. The SeMet C9A *Sp*CstR crystals were obtained in a different space group (P21) and diffracted to ≈2.3 Å. The structure was solved by molecular replacement (MR) using MoRDa^70^ and PDB 3AAI as the search model. Phases were improved using Autosol in PHENIX^71^ and the dataset obtained at 1.77118 nm in space group C2 gave four Sulfur sites with fractional occupancies of 0.81-0.91. A Figure of Merit (FOM) of 0.38 was obtained and an incomplete model with *R*_work_/*R*_free_ of 0.374/0.434 was built. Successive cycles of automatic building in Autobuild (PHENIX) and manual building in Coot,^72^ as well as refinement (PHENIX Refine^73^) led to a complete model. These coordinates were then used as search model for MR to phase datasets of SeMet C9A *Sp*CstR, with four molecules in the asymmetric unit, and native C9A *Sp*CstR and C9A/N55A *Sp*CstR. Some side chains for several residues in these models were not intentionally built due to the lack of electron density. The SeMet C9A *Sp*CstR model obtained at pH 7.5 is characterized by relatively high B-factors (Table S4), very weak electron density in the α1-α2 and α2-α3 loops and a loss of helicity in the α3 helix on one of four chains in the asymmetric unit. In contrast, the C9A/N55A *Sp*CstR model in space group I 2 2 2 shows an *R*_work_/*R*_free_ of 0.1998/0.2251 with excellent geometry and no Ramachandran outliers (Table S4). All crystallographic data acquisition and refinement statistics are shown in Table S4.

### NMR spectroscopy

Uniformly ^15^N, ^13^C, ^2^H protein was expressed in *E. coli* BL21 (DE3) cells, as described above, except in M9 minimal medium containing 1 kg D_2_O, with 1.0 g of ^15^NH_4_Cl and 2 g ^13^C_6_,^2^H-glucose as the sole nitrogen and carbon sources, respectively. Uniformly ^15^N-labeled protein was expressed in M9 minimal medium containing 1 L H_2_O, as well as 1.0 g of ^15^NH_4_Cl as the sole nitrogen source. NMR samples for backbone assignment contained 0.2-0.6 mM protein, with 25 mM MES pH 5.5, 150 mM NaCl, and 10% v/v D_2_O, with 0.3 mM 2,2-dimethyl-2-silapentanesulfonic acid (DSS) as an internal reference. NMR spectra were recorded at 35 °C on a Bruker Avance Neo 600 MHz spectrometer equipped with a cryogenic probe in the METACyt Biomolecular NMR Laboratory at Indiana University, Bloomington. Backbone chemical shifts were assigned for each mutant at pH 5.5 using TROSY versions of the following standard triple-resonance experiments: HNCACB, HNCOCACB, and HNCO^74^ using non-uniform sampling with Poisson gap schedules^75^. Data were collected using Topspin 4.0.7 (Bruker), processed using NMRPipe^76^ and istHMS^77^, and analyzed using CARA^78^ and NMRFAM-Sparky,^79^ all on NMRbox^80^. Assignments at pH 7.5 were obtained by titration from pH 5.5 through pH 6.0 and pH 6.5. Backbone chemical shift perturbations (CSP) were calculated using ^1^H and ^15^N chemical shifts with Δδ=((ΔδH)^2^+ 0.2(ΔδN)^2^)^1/2^. Chemical shift assignments have been deposited at the BMRB under accession codes 50893 and 50894 for *Sp*CstR C9A and *Sp*CstR C9A N55A, respectively.

### Molecular dynamics simulations

Initial coordinates for molecular dynamics simulations for C9A/N55A *Sp*CstR were obtained from reduced crystal structure reported here (PDB *code* 7MQ3). In the case of C9A *Sp*CstR, two sets of initial coordinates were considered: an *in silico* mutation of N55 was performed on the C9A/N55A *Sp*CstR that has both high resolution and resolved N- and C-terminal regions, and the C9A *Sp*CstR crystallographic structure obtained in pH 7.5. In both C9A *Sp*CstR MD simulations, the tetrameric assembly state remains stable throughout the simulations (Fig. S8), however the *in-silico* mutant is of remarkable higher quality and was used for the analysis. Crystallization ions, organic molecules and water molecules were not considered part of the initial coordinates of the system.

All molecular dynamics simulations were performed using the Gromacs MD engine^81^ using the Amber99SB force field.^82^ Initial coordinates of the tetrameric protein structures were solvated in an octahedral box of TIP3P water molecules^83^ and sodium ions to achieve electroneutrality. All simulations were performed using periodic boundary conditions and Ewald sums to treat long range electrostatic interactions, the SHAKE algorithm^84^ to keep bonds involving hydrogen atoms at their equilibrium length, 2 fs time step for the integration of Newton’s equations, and the Berendsen thermostat and barostat to control the system temperature and pressure respectively. Equilibration consisted of an energy minimization of the initial structures, followed by a slow, 2 ns long heating up to 300 K (in the NVT ensemble), and slow relaxation of positional restraints to the protein atoms in the NPT ensemble. Production MD runs consisted of 400 ns long trajectories, but we considered 200 ns of equilibrium runs to ensure a valid comparison between authentic C9A/N55A *Sp*CstR and the *in-silico* C9A *Sp*CstR mutant. Frames were collected at 500 ps intervals, which were subsequently used to analyze the production trajectories. Trajectory analyses were carried out using the CPPTRAJ module of AmberTools^85^ and visualization and figures were generated using VMD^86^ and Pymol (v2.4.0, Schrödinger, LLC).

## Supporting Information

The Supporting Information is available free of charge on the ACS Publications Web site and includes Supplementary Tables S1-S7, Supplementary Figures S1-S10 and Supplementary References.

## Author Information

### Corresponding authors

**Daiana A. Capdevila** – Fundación Instituto Leloir, Av. Patricias Argentinas 435, Buenos Aires C1405BWE, Argentina. https://orcid.org/0000-0002-0500-1016

**David P. Giedroc** – Departments of Chemistry, Indiana University, 800 E. Kirkwood Ave, Bloomington, IN 47405-7102, United States., https://orcid.org/0000-0002-2342-1620

### Authors

**Joseph N. Fakhoury** – Department of Chemistry, Indiana University, 800 E. Kirkwood Ave, Bloomington, IN 47405-7102, United States.

**Yifan Zhang** – Departments of Chemistry and of Molecular and Cellular Biochemistry, Indiana University, 800 E. Kirkwood Ave, Bloomington, IN 47405-7102, United States.

**Katherine A. Edmonds** – Department of Chemistry, Indiana University, 800 E. Kirkwood Ave, Bloomington, IN 47405-7102, United States. https://orcid.org/0000-0002-1282-9858

**Mauro Bringas** – Fundación Instituto Leloir, Av. Patricias Argentinas 435, Buenos Aires C1405BWE, Argentina.

**Justin L. Luebke** – Department of Chemistry, Indiana University, 800 E. Kirkwood Ave, Bloomington, IN 47405-7102, United States.

**Giovanni Gonzalez-Gutierrez** – Department of Molecular and Cellular Biochemistry, Indiana University, Bloomington, IN 47405, United States. https://orcid.org/0000-0002-1044-943X

## Notes

The authors declare no conflicts of interest.

## Acknowledgments

We thank members of the Giedroc and Capdevila laboratories for helpful comments on the manuscript. We gratefully acknowledge use of the Macromolecular Crystallography Facility at the Molecular and Cellular Biochemistry Department, Indiana University, Bloomington. We also thank J. Nix for his assistance during X-ray data collection at beamline 4.2.2, ALS. We thank H. Wu for his help with NMR data acquisition. We thank the INQUIMAE-DQIAyQF cluster (FCEN, UBA) for providing computational resources.

