## Supplementary information for "Functional asymmetry and chemical reactivity of CsoR family persulfide sensors"

This file contains Supplementary Tables S1-S7 and Supplemental Figures S1-S10 and Supplementary References.

This Supporting Information includes:


**Supporting Tables** page

**Table S1.** SSN cluster sizes ranked according to the number of unique sequences… S3
**Table S2.** Summary of observed and calculated monoisotopic masses for cysteine-containing peptides in the rPA-MS experiments……………………………………….. S5
**Table S3.** Summary of pulsed alkylation rate constants……………………………… S6
**Table S4.** Crystallographic data collection and refinement statistics…………………... S7
**Table S5.** Summary of the calculated and observed masses of all oxidation products of intact proteins with ESI-MS used in the kinetic profiling experiments with hydrogen peroxide and tetrathionate……………………………………………………………….. S9
**Table S6.** Summary of the calculated and observed masses of all oxidation products of intact proteins with ESI-MS used in the kinetic profiling experiments with glutathione disulfide…………………………………………………………………….. S10
**Table S7.** Summary of the calculated and observed masses of all oxidation products of intact proteins with ESI-MS used in the kinetic profiling experiments with the cysteine persulfide donor……………………………………………………………………. S12

**Supporting Figures**

**Figure S1.** Genomic neighborhood analysis of all functionally characterized CsoR family members………………………………………………………………………………. S13
**Figure S2.** Sequence similarity network analysis of Ni/Co-sensors and functionally uncharacterized clusters 6-9………………………………………………………………... S15
**Figure S3.** Pulsed alkylation mass spectrometry of the N-terminal Cys residue in *Sa*CstR vs. *Sa*CsoR…………………………………………………………………………. S17
**Figure S4.** Multiple sequence alignments and thermodynamics of DNA operator binding for CstRs studied here……………………………………………………………... S18
**Figure S5.** Small angle x-ray scattering data and analysis obtained for each of the three *Sp*CstR structures……………………………………………………………………. S19
**Figure S6.** $^{15}$N,$^{1}$H TROSY spectra of reduced C9A *Sp*CstR at pH 5.5 and pH 7.5….. S20
**Figure S7.** Solution NMR analysis of C9A/N55A *Sp*CstR vs. the parent C9A *Sp*CstR S21
**Figure S8.** Molecular dynamics simulations of C9A/N55A *Sp*CstR, C9A *Sp*CstR and C9A/G26E *Sp*CstR………………………………………………………………………... S22
**Figure S9.** Representative reaction profiles of CstRs with a 20-fold molar protomer excess of tetrathionate………………………………………………………………………. S23
**Figure S10.** Representative reaction profiles of CstRs with a 20-fold molar protomer excess of cystine……………………………………………………………………………... S24

**Supporting References**……………………………………………………………………. S24

**Table S1.** SSN cluster sizes ranked according to the number of unique sequences[a]

| Cluster Number | UniProt Cluster Size | UniRef90 Cluster Size | Fraction total sequences (cumulative %) | Cluster Number | UniProt Cluster Size | UniRef90 Cluster Size | % total sequences |
|---|---|---|---|---|---|---|---|
| 1 | 9450 | 3563 | 0.2909 | 61 | 22 | 21 | |
| 2 | 6010 | 1082 | 0.1850 | 62 | 24 | 15 | |
| 3 | 3175 | 1115 | 0.0977 | 63 | 22 | 9 | |
| 4 | 1816 | 565 | 0.0559 | 64 | 23 | 8 | |
| 5 | 1836 | 586 | 0.0565 | 65 | 23 | 18 | |
| 6 | 870 | 376 | 0.0268 | 66 | 23 | 17 | |
| 7 | 843 | 361 | 0.0259 | 67 | 20 | 11 | |
| 8 | 800 | 358 | 0.0246 | 68 | 21 | 9 | |
| 9 | 496 | 154 | 0.0153 | 69 | 24 | 6 | |
| 10 | 483 | 106 | 0.0149 (79.3) | 70 | 26 | 5 | 0.0070 (93.9) |
| 11 | 379 | 64 | 0.0117 | 71 | 24 | 18 | |
| 12 | 372 | 162 | 0.0115 | 72 | 24 | 13 | |
| 13 | 343 | 236 | 0.0106 | 73 | 21 | 17 | |
| 14 | 282 | 88 | 0.0086 | 74 | 19 | 11 | |
| 15 | 215 | 161 | 0.0066 | 75 | 19 | 17 | |
| 16 | 172 | 117 | 0.0052 | 76 | 18 | 16 | |
| 17 | 160 | 87 | 0.0049 | 77 | 19 | 4 | |
| 18 | 155 | 47 | 0.0047 | 78 | 17 | 8 | |
| 19 | 132 | 77 | 0.0040 | 79 | 18 | 6 | |
| 20 | 131 | 88 | 0.0040 (86.6) | 80 | 18 | 3 | 0.0061 (94.5) |
| 21 | 118 | 66 | 0.0036 | 81 | 14 | 11 | |
| 22 | 105 | 43 | 0.0032 | 82 | 17 | 16 | |
| 23 | 93 | 46 | 0.0028 | 83 | 16 | 9 | |
| 24 | 97 | 55 | 0.0030 | 84 | 13 | 4 | |
| 25 | 87 | 53 | 0.0027 | 85 | 14 | 9 | |
| 26 | 97 | 8 | 0.0030 | 86 | 14 | 9 | |
| 27 | 85 | 41 | 0.0026 | 87 | 12 | 10 | |
| 28 | 81 | 47 | 0.0025 | 88 | 14 | 12 | |
| 29 | 83 | 3 | 0.0026 | 89 | 14 | 4 | |
| 30 | 72 | 41 | 0.0022 (89.4) | 90 | 10 | 7 | 0.0042 (95.0) |
| 31 | 79 | 59 | | 91 | 12 | 10 | |
| 32 | 62 | 18 | | 92 | 11 | 10 | |
| 33 | 61 | 31 | | 93 | 14 | 14 | |
| 34 | 57 | 29 | | 94 | 13 | 12 | |
| 35 | 51 | 27 | | 95 | 13 | 11 | |
| 36 | 55 | 27 | | 96 | 14 | 9 | |

| | | | | | | | |
|---|---|---|---|---|---|---|---|
| 37 | 53 | 30 | | 97 | 14 | 1 | |
| 38 | 50 | 34 | | 98 | 14 | 1 | |
| 39 | 54 | 12 | | 99 | 13 | 2 | |
| 40 | 54 | 4 | 0.018 (91.2) | 100 | 13 | 8 | 0.0040 (95.4) |
| 41 | 50 | 39 | | 101 | 10 | 8 | |
| 42 | 41 | 16 | | 102 | 12 | 5 | |
| 43 | 42 | 17 | | 103 | 11 | 8 | |
| 44 | 41 | 8 | | 104 | 11 | 11 | |
| 45 | 36 | 23 | | 105 | 12 | 7 | |
| 46 | 35 | 20 | | 106 | 11 | 4 | |
| 47 | 36 | 8 | | 107 | 13 | 10 | |
| 48 | 32 | 24 | | 108 | 9 | 5 | |
| 49 | 32 | 13 | | 109 | 12 | 2 | |
| 50 | 34 | 3 | 0.0117 (92.4) | 110 | 13 | 1 | 0.0035 (95.7) |
| 51 | 31 | 11 | | 111 | 11 | 9 | |
| 52 | 26 | 14 | | 112 | 11 | 9 | |
| 53 | 31 | 6 | | 113 | 12 | 9 | |
| 54 | 29 | 13 | | 114 | 11 | 9 | |
| 55 | 25 | 10 | | 115 | 12 | 3 | |
| 56 | 23 | 9 | | 116 | 11 | 10 | |
| 57 | 25 | 16 | | 117 | 10 | 6 | |
| 58 | 27 | 19 | | 118 | 10 | 6 | |
| 59 | 24 | 12 | | 119 | 8 | 6 | |
| 60 | 26 | 22 | 0.0082 (93.2) | 120 | 10 | 7 | 0.0033 (96.0) |

[a]All clusters (from 511 total non-singleton clusters) with more than 10 Uniref90 sequences are shown here. [b]% total unique sequences of 32,496 sequences (11,726 UniRef90 sequences) for clusters 1-30 are shown, then grouped in groups of 10 clusters for 41-50, 51-60, 61-70, etc.

**Table S2**. Summary of observed and calculated monoisotopic masses for cysteine-containing peptides in the rPA-MS experiments.[a]

| | | | Mass of Cys-Containing Peptides (Da) | | | | |
|---|---|---|---|---|---|---|---|
| **Cys** | **Peptide** | **Sequence** | **Cys-SH** | **Cys-S-$d_5$-NEM** | | **Cys-S-$H_5$-NEM** | |
| | | | Calc. | Calc. | Obs. | Calc. | Obs. |
| *Sp*CstR C9 | 6-12 | (K)YITCLKR(S) | 896.5 | 1126.5 | 1126.6 | 1121.5 | 1121.5 |
| *Sp*CstR C30 | 23-41 | (K)MIEGDRDCADIVT QLTAVR(S) | 2106.0 | 2236.1 | 2236.1 | 2231.1 | 2231.1 |
| *Sp*CstR C59 | 47-73 | (R)VIEMIITENLTECIN QPLDDSEAQKER(L) | 3131.5 | 3261.5 | 3261.4 | 3256.6 | 3256.6 |
| *Sm*CstR C30 | 23-41 | (K)MIEEDRDCADIVT QLTAVK(S) | 2150.0 | 2280.1 | 2280.1 | 2275.1 | 2275.1 |
| *Sm*CstR C59 | 47-73 | (R)VIEMIITENLTECIN QPLDDPEAQKER(L) | 3141.5 | 3271.6 | 3271.6 | 3266.6 | 3266.5 |
| *Ef*CstR C31 | 30-42 | (K)ECIDVITQLSA VR(S) | 1446.8 | 1576.8 | 1576.8 | 1571.8 | 1571.8 |
| *Ef*CstR C60 | 59-70 | (K)HCFENPEKDPK(E) | 1343.6 | 1473.7 | 1473.6 | 1468.7 | 1468.6 |
| *Sa*CstR C31 | 24-42 | (K)MMEEGKD**C**KDVIT QISASK(S) | 2113.0 | 2243.1 | 2243.1 | 2238.0 | 2238.0 |
| *Sa*CstR C60 | 48-62 | (R)LMGIIISENLIE**C**VK (A) | 1674.9 | 1805.0 | 1804.9 | 1800.0 | 1799.9 |
| *Sa*CsoR C41 | 34-49 | (R)MIEEDVY**C**DDVLT QIR(A) | 1941.9 | 2072.0 | 2072.0 | 2066.9 | 2066.9 |
| *Sa*CsoR C70 | 69-93 | (K)S**C**IMNKVNQGAQE EAMEELLVTFQK(L) | 2840.4 | 2970.3 | 2970.4 | 2965.4 | 2965.5 |

[a]Cysteines were alkylated with either $H_5$-NEM or $d_5$-NEM followed by tryptic digestion with peptides detected by MALDI-TOF mass spectrometry (see **Fig. 2**, main text).

**Table S3**. Summary of pulsed alkylation rate constants

| Protein | N-terminal Cys<br>$k$ ($M^{-1}$ $min^{-1}$) | C-terminal Cys<br>$k_{fast}$ ($M^{-1}$ $min^{-1}$) ($A^b$) | $k_{slow}$ ($M^{-1}$ $min^{-1}$) (A) |
|---|---|---|---|
| C9A *Sp*CstR | 4.0 (±0.6) x$10^4$ | 3.2 (±1.3) x$10^4$ (0.20 ±0.05) | 1600 (±200) (0.80) |
| *Sm*CstR | 9.8 (±0.7) x$10^4$ | 7.3 (±3.6) x$10^4$ (0.25 ±0.05) | 3800 (±400) (0.75) |
| *Ef*CstR | 1.22 (±0.03) x$10^4$ | 8100 (±900) | n.d.[c] |
| *Sa*CstR | 2100 (± 200) | 8200 (±800) | 610 (±170) |
| *Sa*CsoR[a] | *44 (±1.2)* | 2.2 (±0.4) x$10^4$ | 200 (±50) |

[a]The more N-terminal and C-terminal cysteine residues for *Sa*CsoR are C41 and C70, respectively, with other Cys residue numbers in the other CstRs shown in Table S2. [b]A, amplitude. [c]Results of a single-exponential fit are shown.

**Table S4.** Crystallographic data collection and refinement statistics[a]

| | C9A *Sp*CstR (pH 5.5) | C9A *Sp*CstR (pH 5.5) | SeMet C9A *Sp*CstR (pH 7.5) | C9A/N55A *Sp*CstR (pH 6.0) |
|---|---|---|---|---|
| *Data collection* | | | | |
| Wavelength (Å) | 1.00003 | 1.77118 | 0.97625 | 0.97625 |
| Space group | C2 | C2 | P 21 1 | I 2 2 2 |
| *Cell dimensions* | | | | |
| a, b, c (Å) | 93.30 54.87 57.51 | 93.84 55.14 57.65 | 59.44 45.57 64.95 | 31.91 66.93 76.05 |
| $\alpha, \beta, \gamma$ (°) | 90.00 115.73 90.00 | 90.00 115.71 90.00 | 90.00 112.85 90.00 | 90.00 90.00 90.00 |
| Resolution (Å) | 43.04 – 2.02 (2.07 – 2.02) | 31.05 – 2.02 (2.07 – 2.02) | 45.57 – 2.29 (2.37 – 2.29) | 29.42 – 1.40 (1.42 – 1.40) |
| $R_{sym}$ | 0.047 (0.759) | 0.063 (1.229) | 0.069 (0.925) | 0.047 (1.125) |
| $R_{meas}$ | 0.055 (0.910) | 0.065 (1.283) | 0.074 (0.988) | 0.049 (1.223) |
| $R_{pim}$ | 0.029 (0.497) | 0.018 (0.364) | 0.028 (0.374) | 0.014 (0.474) |
| Total reflections | 61538 (4158) | 228856 (15517) | 103023 (9921) | 172088 (5319) |
| No. unique reflections | 17243 (1273) | 16923 (1274) | 14665 (1408) | 16441 (814) |
| CC1/2 | 0.999 (0.633) | 0.999 (0.840) | 0.999 (0.811) | 0.999 (0.785) |
| $I/\sigma(I)$ | 10.8 (1.1) | 31.5 (1.9) | 16.2 (1.7) | 27.4 (1.5) |
| Completeness (%) | 99.6 (99.9) | 96.6 (99.9) | 99.9 (100.0) | 99.8 (100.0) |
| Multiplicity | 3.6 (3.3) | 13.5 (12.2) | 7.0 (7.0) | 10.5 (6.5) |
| Wilson B-factor | 36.70 | 35.35 | 48.39 | 18.87 |
| *Refinement* | | | | |
| Resolution (Å) | 43.04 – 2.02 (2.15 – 2.02) | | 36.26 – 2.29 (2.36 – 2.29) | 28.80 – 1.40 (1.45 – 1.40) |
| No. unique reflections | 17236 (1731) | | 14654 (1435) | 16367 (1616) |
| $R_{work}$ | 0.2048 (0.2791) | | 0.2589 (0.3268) | 0.1998 (0.2986) |
| $R_{free}$ | 0.2576 (0.3785) | | 0.2991 (0.3442) | 0.2251 (0.3458) |
| CC(work) | 0.966 (0.812) | | 0.959 (0.804) | 0.963 (0.878) |
| CC(free) | 0.942 (0.677) | | 0.958 (0.687) | 0.982 (0.710) |
| *R.m.s.d values* | | | | |
| Bond lengths (Å) | 0.008 | | 0.015 | 0.004 |
| Bond angles (°) | 0.978 | | 1.75 | 0.660 |
| *No. atoms* | | | | |

| | | | |
|---|---|---|---|
| Protein | 1931 | 2304 | 671 |
| Ligand/ions | 41 | | 13 |
| solvent | 116 | 21 | 95 |
| *B-factors* (Å$^2$) | | | |
| Protein | 52.71 | 75.28 | 33.24 |
| ligand/ions | 68.51 | | 45.94 |
| solvent | 53.31 | 70.13 | 43.32 |
| *Ramachandran plot* | | | |
| Favored (%) | 98.75 | 97.86 | 100 |
| Allowed (%) | 1.25 | 2.14 | 0 |
| Clashscore | 6.53 | 3.23 | 5.05 |
| Rotamer outliers (%) | 0.0 | 0.0 | 0.0 |
| *PDB code* | 7MQ1 | 7MQ2 | 7MQ3 |

[a]Highest-resolution shell values are shown in parentheses.

**Table S5.** Summary of the calculated and observed masses of all oxidation products of intact proteins with ESI-MS used in the kinetic profiling experiments with hydrogen peroxide (refer to Fig. 6) and tetrathionate (refer to Fig. S8).

| | | C9A *Sp*CstR | | C9A/N55A *Sp*CstR | | *Sm*CstR | |
|---|---|---|---|---|---|---|---|
| $H_2O_2$ species symbol | Structure schematic | Expected mass (Da) | Observed mass (Da) | Expected mass (Da) | Observed mass (Da) | Expected mass (Da) | Observed mass (Da) |
| 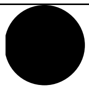 | 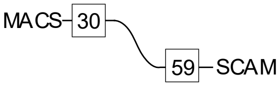  | 9837 | 9838 | 9794 | 9795 | 10236 | 10236 |
| 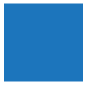 | 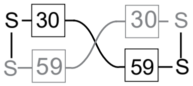  | 19442 | 19443 | 19356 | 19356 | 20240 | 20239 |
| 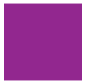 | 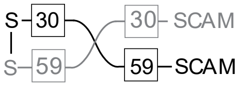  | 19556 | 19558 | 19473 | 19473 | 20354 | 20354 |
| Tetrathionate species symbol | | | | | | | |
| 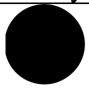 | 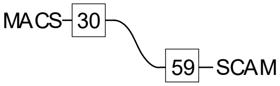  | 9837 | 9838 | 9794 | 9796 | 10236 | 10236 |
| 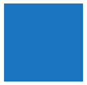 | 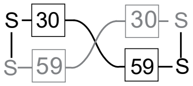  | 19442 | 19443 | 19356 | 19358 | 20240 | 20239 |

**Table S6.** Summary of the calculated and observed masses of all oxidation products of intact proteins with ESI-MS used in the kinetic profiling experiments with glutathione disulfide (GSSG) (refer to Fig. S10).

| | | C9A *Sp*CstR | | C9A/N55A *Sp*CstR | |
|---|---|---|---|---|---|
| GSSG species symbol | Structure schematic[a] | Expected mass (Da) | Observed mass (Da) | Expected mass (Da) | Observed mass (Da) |
| 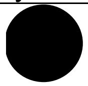 | 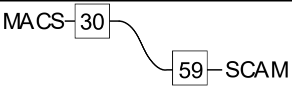  | 9837 | 9838 | 9794 | 9795 |
| 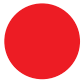 | 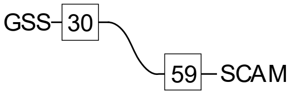  | 10086 | 10086 | 10043 | 10044 |
| 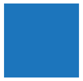 | 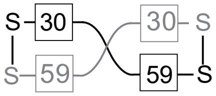  | 19442 | 19444 | 19356 | 19358 |
| 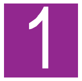  | 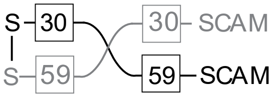  | 19556 | 19559 | 19470 | 19474 |
| 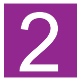  | 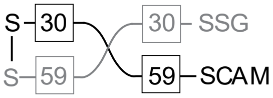  | 19805 | 19808 | 19719 | 19723 |
| 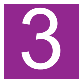  | 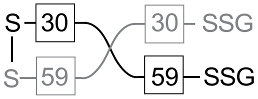  | 20054 | 20057 | 19968 | 19971 |

[a]All structures are drawn to show addition to the attacking Cys (C30 in *Sp*CstR and *Sm*CstR, C31 in *Ef*CstR) but may be a mixture of addition to both C30 and C59. G, glutathione

**Table S7.** Summary of the calculated and observed masses of all oxidation products of intact proteins with ESI-MS used in the kinetic profiling experiments with the cysteine persulfide (CysSSH) donor (refer to Fig. 7).

| | | C9A *Sp*CstR | | C9A/N55A *Sp*CstR | | *Sm*CstR | | *Ef*CstR | |
|---|---|---|---|---|---|---|---|---|---|
| CysSSH species symbol | Structure schematic[a] | Expected mass (Da) | Observed mass (Da) | Expected mass (Da) | Observed mass (Da) | Expected mass (Da) | Observed mass (Da) | Expected mass (Da) | Observed mass (Da) |
| ● | 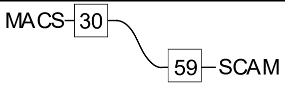  | 9837 | 9837 | 9794 | 9794 | 10236 | 10235 | 10081 | 10082 |
| 1 | 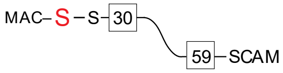  | 9869 | 9869 | 9826 | 9826 | 10268 | 10267 | 10113 | 10114 |
| 2 | 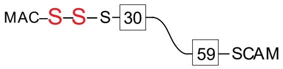  | 9901 | 9900 | 9858 | 9857 | 10300 | 10299 | 10145 | 10145 |
| 3 | 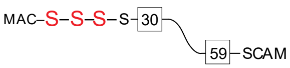  | 9933 | 9931 | 9890 | 9888 | 10332 | 10330 | 10177 | 10175 |
| ■ | 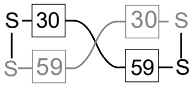  | 19442 | 19442 | 19356 | 19355 | 20240 | n.d.[b] | 19930 | 19930 |
| 1 | 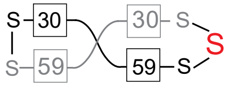  | 19474 | 19473 | 19388 | 19387 | 20272 | n.d. | 19962 | 19963 |
| 2 | 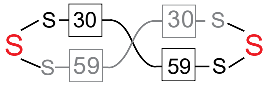  | 19506 | 19506 | 19420 | 19420 | 20304 | n.d. | 19994 | 19995 |
| 3 | 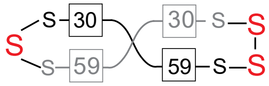  | 19538 | 19538 | 19452 | 19452 | 20336 | n.d. | 20026 | n.d. |
| 4 | 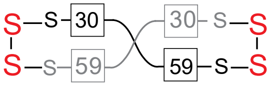  | 19570 | n.d. | 19484 | 19484 | 20368 | n.d. | 20058 | n.d. |

| | | | | | | | | | |
|---|---|---|---|---|---|---|---|---|---|
| 1 | 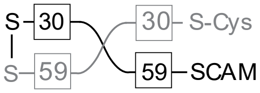  | 19620 | 19622 | 19534 | 19536 | 20418 | 20418 | 20140 | 20142 |
| 2 | 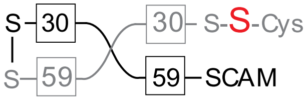  | 19652 | 19653 | 19566 | 19567 | 20450 | 20450 | 20172 | 20172 |
| 3 | 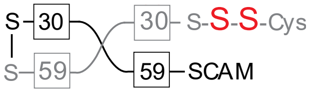  | 19684 | 19685 | 19598 | 19600 | 20482 | 20482 | 20268 | 20269 |

[a]All structures are drawn to show addition to the attacking Cys (C30 in *Sp*CstR and *Sm*CstR, C31 in *Ef*CstR) but may be a mixture of addition to both C30 and C59. [b]not detected

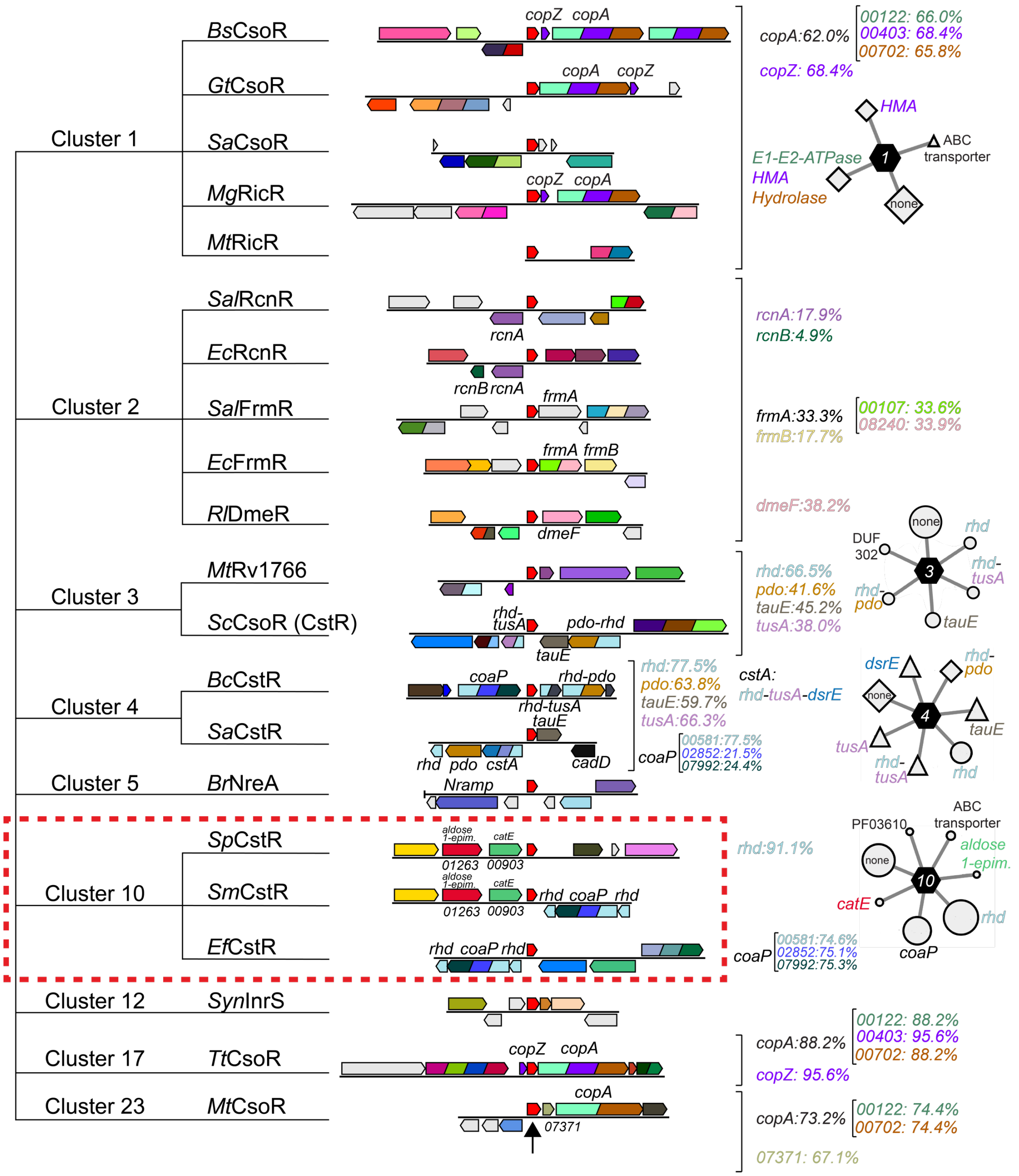


**Figure S1.** Representative Genomic Neighborhood Network (GNN) analyses (±5 genes) on either side of the gene encoding the CsoR family member of interest (*red* gene, marked by the *black* arrow, bottom) of all functionally characterized CsoR family members associated with the indicated SSN cluster (see **Fig. 1**, main text). The SSN cluster 10, boxed in *red*, shows the genomic neighborhood of genes encoding the three CstRs studied here. The proteins encoded by labeled genes are all previously characterized and are indicated by their trivial names and/or Pfam number PF**vwxyz**, *i.e.,* 00122 corresponds to PF00122. *Right*, indicated Pfam domains associated with each trivial gene name, and the percentage with which this Pfam domains appears in all genomic neighborhoods associated with each sequence in the indicated cluster. Spoke-and-wheel diagrams provide a graphical representation of the degree to which an indicated gene co-occurs with the *csoR* family gene associated with cluster 1, 3, 4 and 10 sequences (co-occurrence frequencies ≥20%, ±5 genes). The larger the symbol, the larger the co-occurrence frequency. *copA*, Cu(I)-transporting $P_{1B}$-type ATPase; *copZ*, Cu(I) metallochaperone; HMA, heavy metal ATPase; *rcnAB*, Ni(II)/Co(II) exporter; *frmAB*, formaldehyde detoxification enzymes; *dmeF*, cation diffusion facilitator Ni(II) effluxer; *rhd*, rhodanese sulfurtransferase; *pdo*, persulfide dioxygenase; *tauE*, putative sulfite effluxer; *tusA*, thiouridine sulfurtransferase;, *dsrE*, dissimilatory sulfite reductase sulfurtransferase; *cstA*, multidomain sulfurtransferase[1]; *coaP*, coenzyme A persulfide reductase[2]; *catE*, catechol dioxygenase. See main text for additional details.

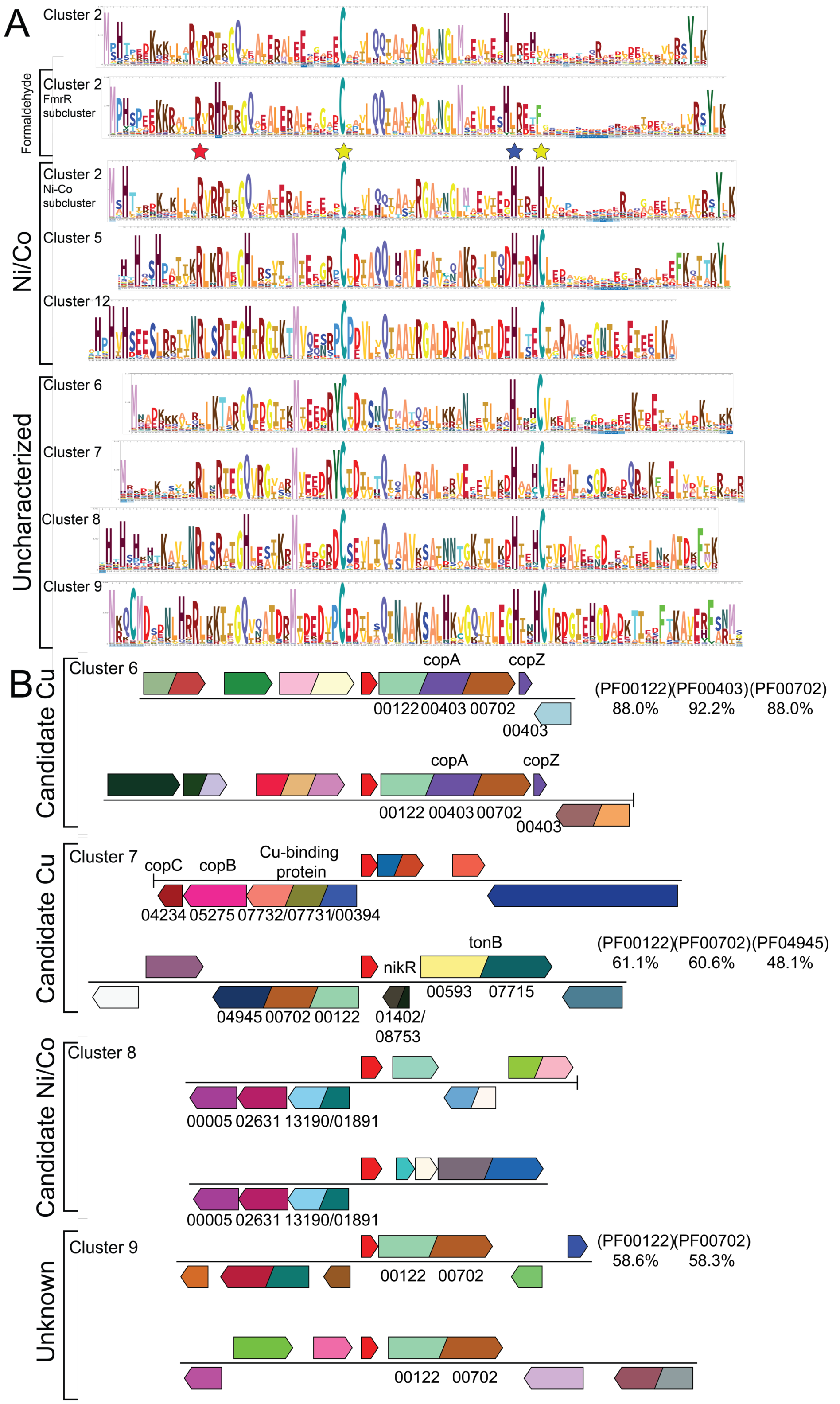


**Figure S2.** Sequence similarity network (SSN) analysis of Ni/Co-sensors and functionally uncharacterized SSN clusters 6-9 (see **Fig. 1**, main text). (A) WebLogo representations of sequence conservation of cluster 2 RcnR/FrmRs (showing both a consensus sequence across the entire cluster of sequences, and for RncRs and FrmRs individually), cluster 5 (NrcB, NreA) and cluster 12 (cyanobacterial InrS) (*top*) and four largest functionally uncharacterized clusters 6-9. The conserved Cys and His residues found in the Cu sensors are marked by *yellow* and *blue* stars respectively, while the *red* star marks the site of the C9A substitution in *Sp*CstR. (B) Representative genomic neighborhood network (GNN) analyses obtained for CsoR family members derived from uncharacterized clusters 6-9. Pfam numbers are shown as described in Fig. S1, with the percentages with which each Pfam domain of *copA* appears in genomic neighborhoods associated with each sequence in the indicated cluster (± five genes on either side of the *csoR* family gene). This analysis suggests that clusters 6 and 7 may well be Cu(I) sensors, cluster 8 may be a Ni/Co sensor, given the propensity of His residues clustered in the N-terminal region (like found in cluster 12 InrS[3]), while the ligand specificity of cluster 9 cannot be predicted from this analysis alone, but may correspond to a novel CsoR family Cd/Zn sensor. Genes of interest, which are biochemically characterized are labeled with their corresponding pfam number. Representative percentage for are labeled in three clusters that may play a role in Cu(I)-binding.

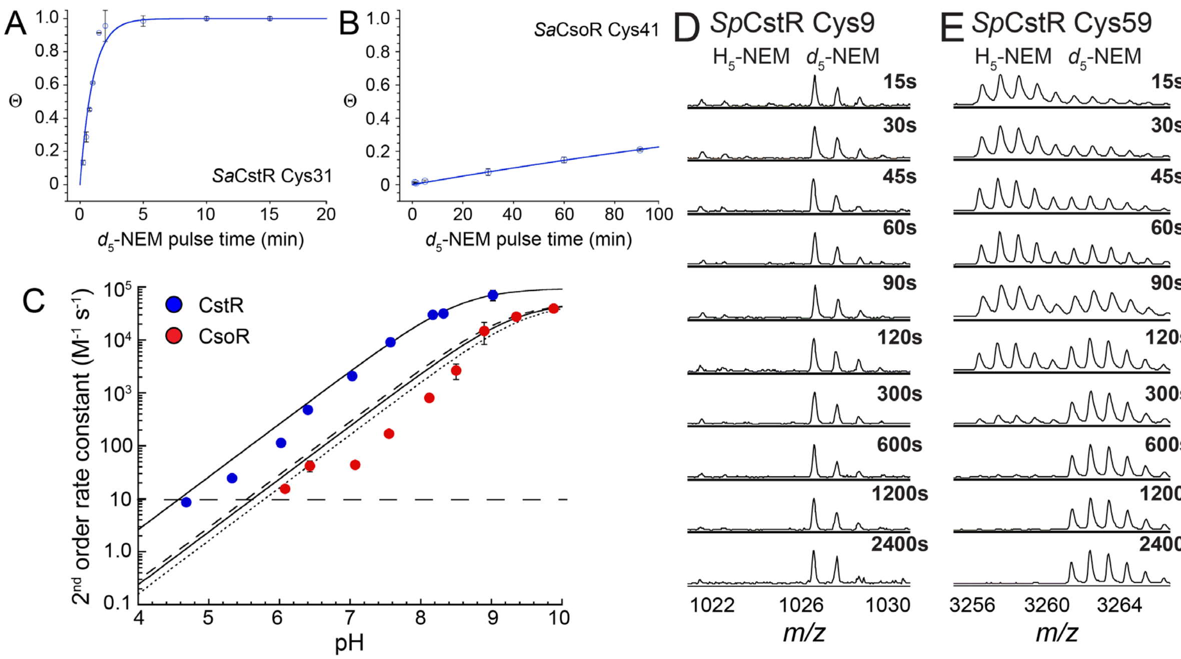


**Figure S3.** Pulsed alkylation mass spectrometry of the N-terminal Cys residue in *Sa*CstR vs. *Sa*CsoR. (A-B) Mol fraction (Θ) of $d_5$-NEM labeled peptide as determined by integration of the $H_5$ and $d_5$-NEM peak areas (see **Fig. 2**, main text). In *Sa*CsoR, Cys41 and Cys70 and analogous to *Sa*CstR Cys31 and Cys60, respectively. Cys31 in *Sa*CstR is analogous to C30 in *Sp*CstR (Fig. S4A). Conditions: 10 mM HEPES, 200 mM NaCl, pH 7.0, anaerobic. (C) Observed p$K_a$ of Cys41 of *Sa*CsoR is approximately 1 pH unit higher than Cys31 of *Sa*CstR. Summary of second order rate constants obtained for Cys31 of *Sa*CstR for the pH range of 4.5-9.0 and Cy41 of *Sa*CsoR between 6.0 and 10.0. The apparent p$K_a$ is ≈8.6±0.1 for Cys31 of *Sa*CstR and 9.3±0.1 for Cys41 of *Sa*CsoR (*solid* line). Simulations of a Cys41 p$K_a$ of 9.25 (*dashed*) and 9.5 (*dotted*) are also shown. Error bars represent the standard deviation from at least three independent experiments. The horizontal dashed line represents the lower limit of reactivity that can be determined from the experimental protocol used here. Selected regions of a series of MALDI-TOF mass spectra obtained by rPA-MS of (D) C9 in wild-type *Sp*CstR and (E) C59 in wild-type *Sp*CstR. Refer to **Fig. 2**, main text for additional details.

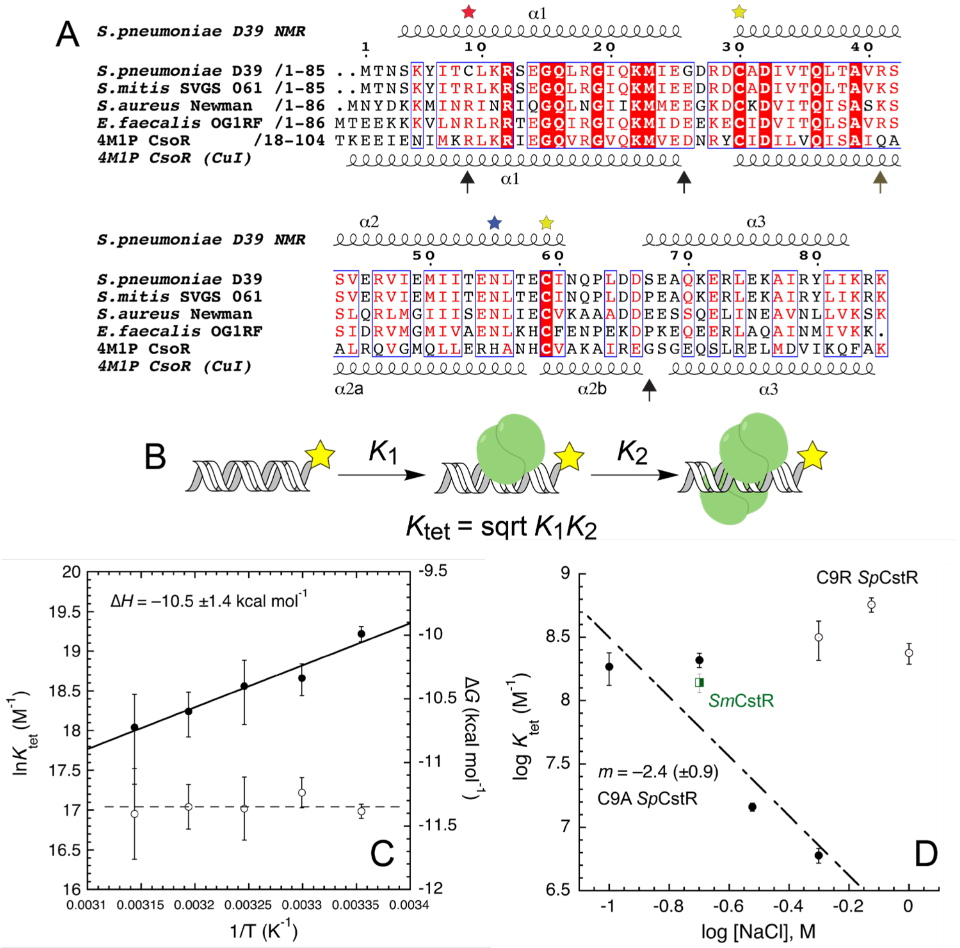


**Figure S4.** Multiple sequence alignments and thermodynamics of DNA operator binding for CstRs studied here. (A) Alignment of three cluster 10 CstRs (*Sp*, *Sm*, *Ef)*, a cluster 4 *Sa*CstR[4] and the structurally characterized cluster 1 *Gt*CsoR (pdb 4M1P).[5] The Cu(I)-coordinating residues are indicated by the *yellow* and *blue* stars, with the position of C9 in *Sp*CstR also indicated (*red* star). The vertical arrows mark the four amino acid differences between *Sp*CstR and *Sm*CstR. The secondary structures for C9A *Sp*CstR and *Gt*CsoR defined by NMR (see Fig. S5 below) and crystallographic data are shown above and below the alignment.[5-6] (B) Schematic representation of the fluorescence anisotropy-based DNA binding assay used here, with the binding model. *Yellow* star, fluorescein dye. (C) van't Hoff plot of the C9A *Sp*CstR *Sm cst* DNA operator binding equilibria. $K_{tet}$, *filled* circles; $\Delta G$, *open* circles. The continuous line through the data are the results of a fit to a van't Hoff model, with $\Delta H$ = –10.5 kcal mol$^{-1}$ and a relatively constant $-T\Delta S$ = ≈ –0.8 kcal mol$^{-1}$ K$^{-1}$. (D) DNA binding by CstR is characterized by a modest [NaCl]-dependence, which according to polyelectrolyte theory reveals the release of ≈2.4 $Na^+$ released or ≈2-3 electrostatic interactions per tetramer.[7] The C9R substitution results in significantly higher affinity for the DNA under the same solution conditions, as anticipated from previous findings in Cu(I)-sensing CsoRs.[8-10]

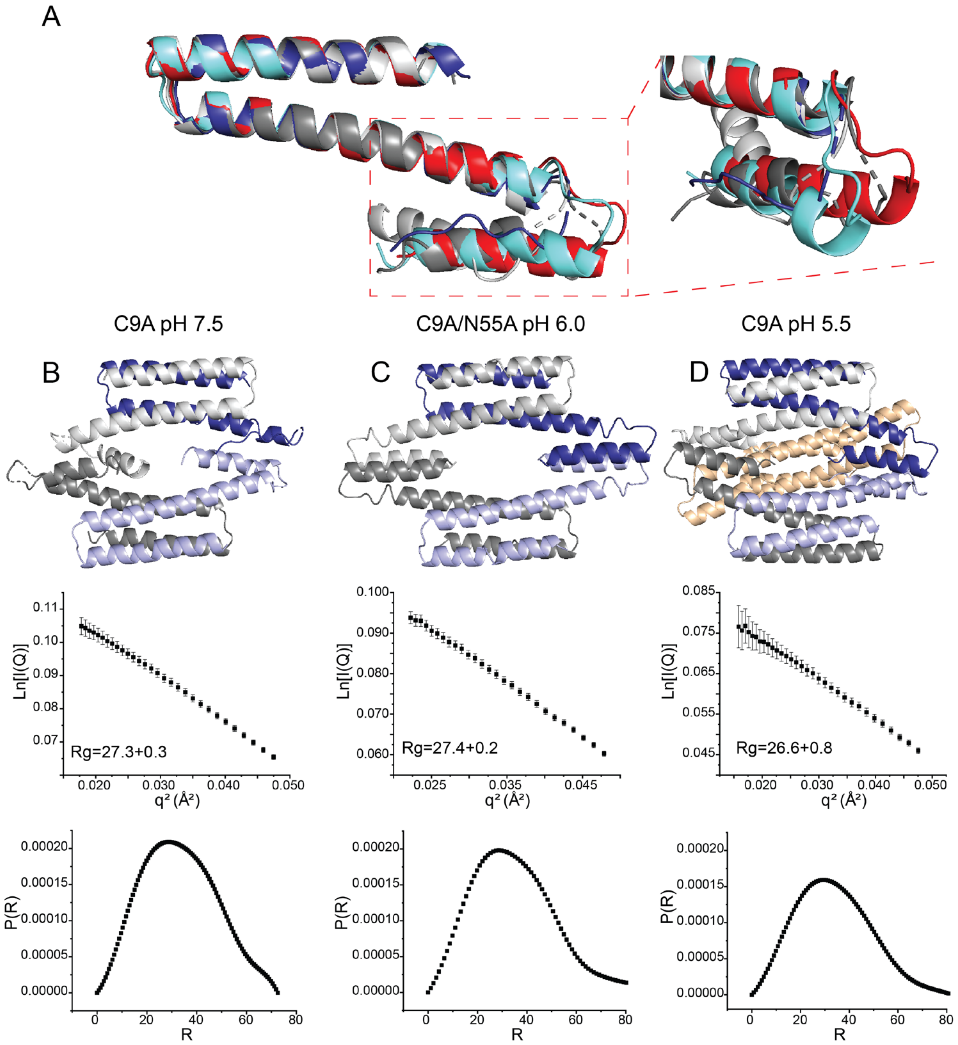


**Figure S5.** (A) Ribbon representation of the crystallographic structure of homotetrameric C9A *Sp*CstR at pH 7.5. A superposition of each of the four chains in the asymmetric unit of the pH 7.5 structure is shown (shaded *white*, *red*, *cyan* and *grey*). See Table S4 for structure statistics. Small angle x-ray scattering data and analysis obtained for the conditions of each of the three *Sp*CstR structures solved here. (B) C9A *Sp*CstR, pH 7.5; (C) C9A *Sp*CstR, pH 5.5; (D) C9A/N55A *Sp*CstR, pH 6.0. *Top*, ribbon structure representations of each structure; *middle*, Guinier plots with the $R_g$ derived from a linear analysis of these data; and *bottom*, Pair distance distribution function (PDDF) plots of each protein were obtained using the distance distribution analysis tool on SAS Data Analysis (primus).[11] These data reveal that the assembly state and hydrodynamic properties of reduced C9A and C9A/N55A *Sp*CstRs do not strongly differ from one another in solution.

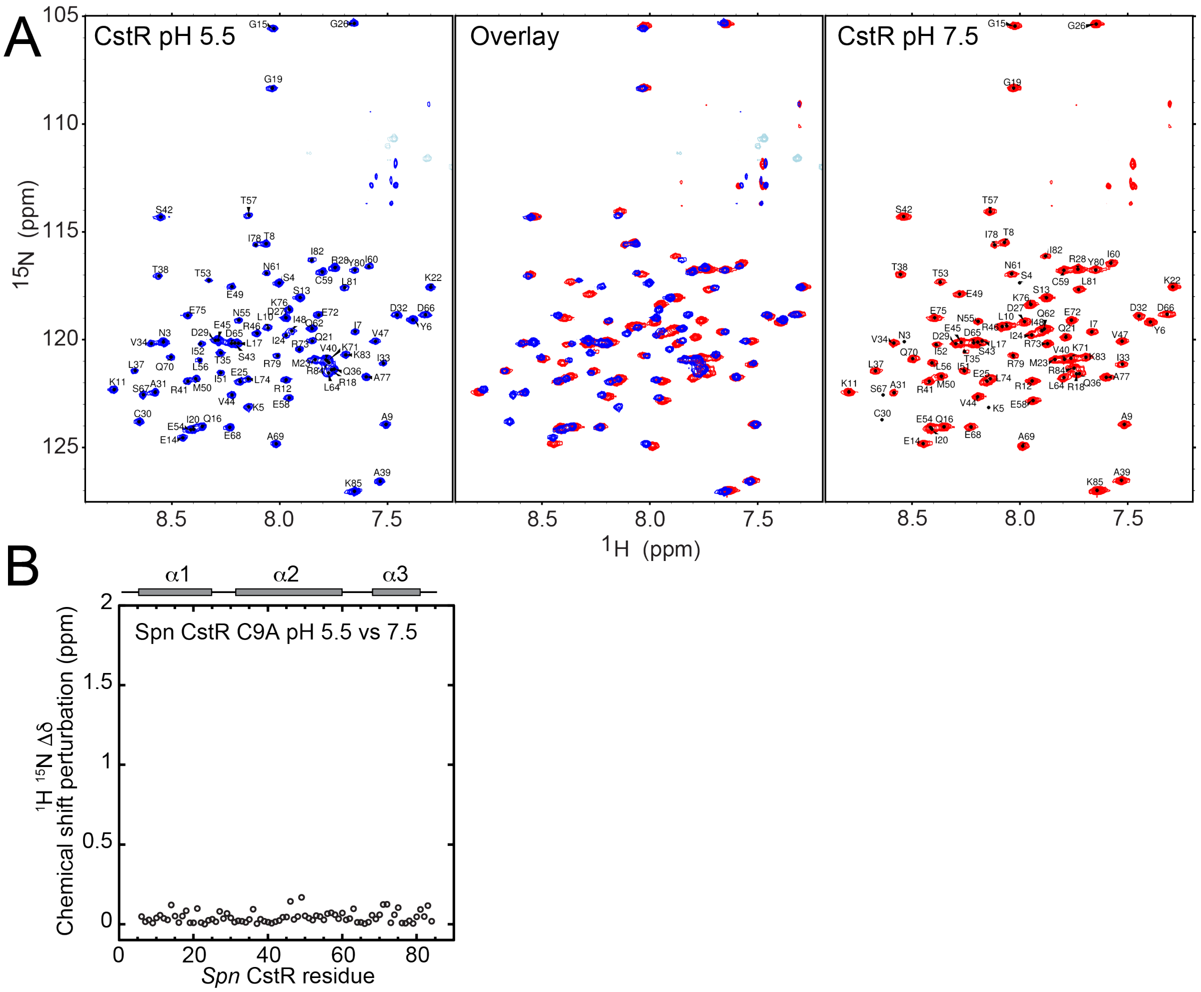


**Figure S6** (A) $^{15}$N,$^{1}$H TROSY spectra of reduced C9A *Sp*CstR at pH 5.5 (*blue* contours) (*left*) compared to the same spectrum of C9A *Sp*CstR at pH 7.5 (*red* contours) (*right*), with both spectra overlaid (*middle*), all acquired at 35 ºC. (B) Chemical shift perturbations of backbone $^{15}$N and the $^{1}H_N$ resonances at pH 5.5 relative to pH 7.5.

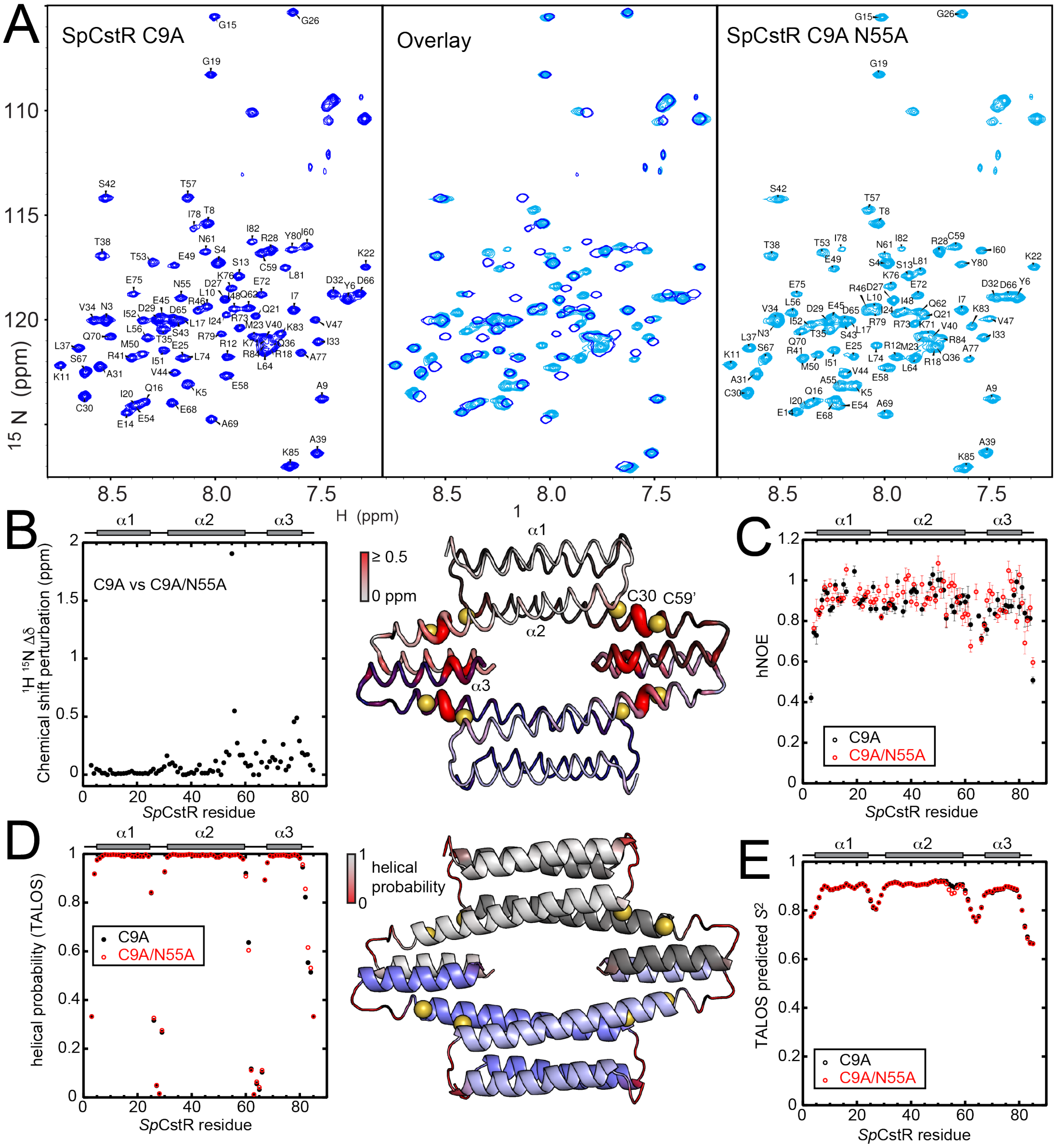


**Figure S7.** Solution NMR analysis of C9A/N55A *Sp*CstR vs. the parent C9A *Sp*CstR. (A) $^{15}$N,$^{1}$H TROSY spectra of reduced C9A *Sp*CstR at pH 5.5 (*blue* contours) (*left*) compared to the same spectrum of C9A N55A *Sp*CstR at pH 5.5 (*cyan* contours) (*right*), with both spectra overlaid (middle), all acquired at 35 ºC. (B) $^{15}$N and $^{1}H_N$ chemical shift perturbation vs. residue number (left), with these changes painted onto the structure of C9A/N55A *Sp*CstR (*right*). (C) Experimentally determined backbone heteronuclear $^{1}$H-$^{15}$N NOE for C9A/N55A (*red*) vs. the parent C9A *Sp*CstR (*black*). The loops that connect the three α-helices as well as the N- and C-terminal tails are more mobile on the sub-ns timescale, with the two variants difficult to distinguish from one another. (D) A comparison of the predicted helical propensities (range 0, no propensity; 1, high propensity) of C9A/N55A (*red*) vs. the parent C9A *Sp*CstR (*black*), based on chemical shifts analysis by TALOS,[12] painted onto the structure of C9A/N55A *Sp*CstR (*right*). (E) TALOS-predicted backbone order parameter ($S^2$) for C9A/N55A (*red*) vs. the parent C9A *Sp*CstR (*black*).

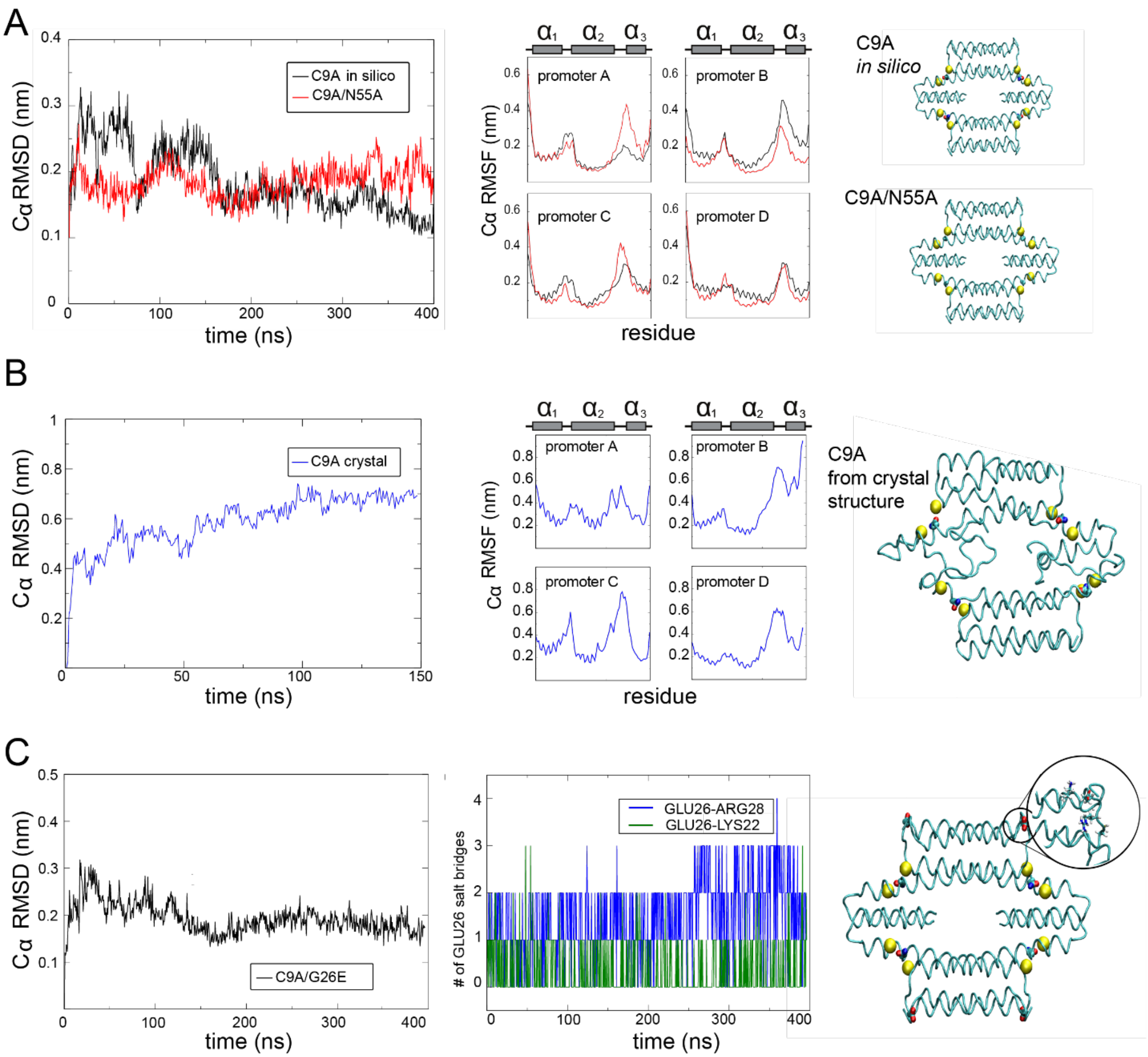


**Figure S8.** Molecular dynamics simulations of C9A/N55A *Sp*CstR, C9A *Sp*CstR and C9A/G26E *Sp*CstR. (A) Cα root mean square deviations (RMSD) as a function of time (*left*), residue number (*middle*) and representative cartoon structures with the $S^{\gamma}$ depicted as spheres (*right*) for the *in silico* C9A *Sp*CstR (*black*) and C9A/N55A *Sp*CstR (*red*) molecular dynamics trajectories. (B) Cα RMSD as a function of time (*left*), residue number (*middle*) and representative cartoon structures with the $S^{\gamma}$ depicted as spheres (*right*) for C9A *Sp*CstR molecular dynamics trajectories using the crystallographic coordinates at pH 7.5 as the initial coordinates. (C) Cα RMSD as a function of time (*left*) for the in C9A/G26E *Sp*CstR (*Sm*CstR-like mutant) molecular dynamics trajectories using the crystallographic initial coordinates from the C9A/N55A *Sp*CstR structure. Time-dependence of the number of salt bridges formed by E26 with K22 (*green*) and R28 (*blue*) of the same protomer (*middle*). Additionally, we observed that there also exists water-mediated interactions between E26 and both basic sidechains, which may contribute to a conformational restriction of the α1-α2 loop. Cartoon indicating the location of E26 in the protein tetrameric structure, with an expanded region that shows basic residues involved in salt bridge formation (*right*).

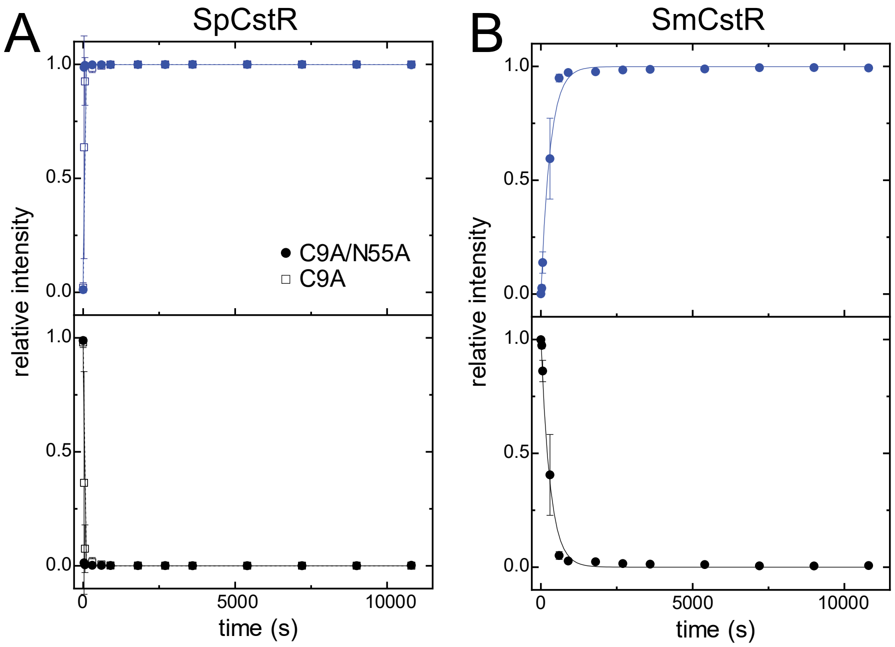


**Figure S9.** Representative reaction profiles of CstRs (60 µM protomer; 15 µM tetramer) with a 20-fold molar protomer excess of tetrathionate. (A) C9A *Sp*CstR (*open* symbols) C9A/N55A *Sp*CstR (*closed* symbols), (B) *Sm*CstR. Peak areas of the reduced monomer (*black* symbols) and disulfide-crosslinked dimer (*blue* circles) units of CstR were quantified as a function of incubation time and plotted as mol fraction of monomer and dimer species. The continuous line through the data represent a simultaneous fit to a first order reaction, with parameters compiled in **Table 2**, main text. *Sm*CstR forms the disulfide at a significantly slower rate relative to the *Sp*CstR derivatives.

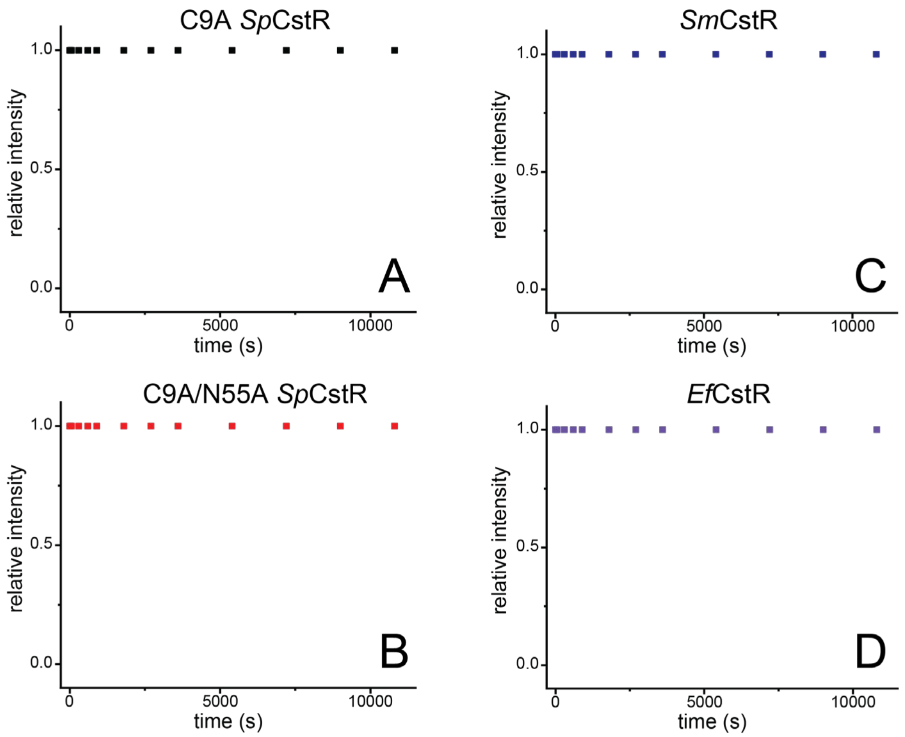


**Figure S10.** Representative reaction profiles of CstRs (60 μM protomer; 15 μM tetramer) with a 20-fold molar protomer excess of cystine. (A) C9A *Sp*CstR, (B) C9A/N55A *Sp*CstR, (C) *Sm*CstR, (D) *Ef*CstR. The only species observed in these reactions is the starting material, reduced monomer.
